# Confidence-supported label-free metabolic imaging with FPhaS fixed phase–autofluorescence microscopy

**DOI:** 10.64898/2026.06.12.731968

**Authors:** Hongming Fan, Jindou Shi, Zhongtian Yang, Alexander Ho, Lingxiao Yang, Kevin K. D. Tan, Edita Aksamitiene, Stephen A. Boppart

**Author notes:** Corresponding author: Stephen A. Boppart, M.D., Ph.D, Beckman Institute for Advanced Science and Technology, University of Illinois Urbana-Champaign, 405 N. Mathews Avenue, Urbana, IL 61801, USA. These authors contributed equally (Hongming Fan, Jindou Shi, Zhongtian Yang). Author emails: H.F.; J.S.; Z.Y.; A.H.; L.Y.; K.K.D.T.; E.A.;.

## Abstract

Label-free optical redox imaging utilizes endogenous NAD(P)H and FAD autofluorescence to evaluate metabolism in living specimens. The conventional optical redox ratio collapses these two channels into a single value; however, it does not indicate whether a pixel has sufficient photon support or the cellular context necessary for quantitative aggregation. To address this limitation, we introduce FPhaS, a fixed-calibration phase– autofluorescence framework that integrates quantitative phase imaging (QPI) with simultaneous label-free autofluorescence multi-harmonic microscopy (SLAM), using fluorescence lifetime imaging (FLIM) solely for validation. Because QPI and SLAM are acquired with the same objective, a unified non-biological calibration aligns phase-derived structural data with the autofluorescence frame, yielding a residual error of 0.39 pixels. This calibration is maintained across all biological specimens. This shared geometric reference enables local evaluation of structural and metabolic information, rather than comparing approximately aligned images. FPhaS decomposes the data into cell presence, ratio credibility, and confidence-supported pooling. We validated FPhaS on A549 cells under high and low-photon conditions; the framework is designed to generalize to other cell and tissue types. Confidence-weighted intensity redox estimates were compared with lifetime-derived measurements within mask-locked cellular regions. Concordance improved exclusively when both the denominator photon support and an independent structural criterion were satisfied. The same reference layer generated cell-level descriptors of metabolic content, metabolic–structural organization, and measurement reliability, while also constraining the CombinedWLS reconstruction under diminished fluorescence acquisition. FPhaS redefines label-free metabolic imaging from producing comprehensive ratio maps to identifying regions where optical evidence substantiates quantitative inference.

## Introduction

In living cells, endogenous autofluorescence from NAD(P)H and FAD has long served as a convenient, label-free indicator of metabolic status^1–4^. The simplest iteration of this measurement combines the two intensity channels into the optical redox ratio, conventionally defined as the FAD signal divided by the sum of the FAD and NAD(P)H signals^5–7^. This simplification holds substantial value. It condenses a complex intracellular metabolic state into a scalar that can be pooled across cells, fields, and conditions. Intensity redox imaging and FLIM capture related but distinct facets of metabolism, namely fluorophore abundance versus enzyme-binding state, so the two are complementary rather than interchangeable^4,7^. The intensity ratio nonetheless remains attractive as a fast, high-throughput readout that scales across cells, fields, and conditions. Although FLIM instrumentation continues to advance toward higher speed and three-dimensional operation^8^, lifetime acquisition still carries a higher photon-count and throughput cost when used as the primary modality at scale, which is the regime FPhaS targets.

Calculating the ratio is not the primary challenge. The real difficulty lies in the excessive permissiveness of the computation. A redox ratio will be computed for any pixel where both autofluorescence channels have values; however, this metric does not address critical concerns such as the strength of the denominator, the reliability of the pixel as part of the cellular interior, or the effects of boundary mixing and background correction that may render the estimate fragile. When such pixels are combined, weak measurement support can be mistaken for biological heterogeneity^3,9^. Although thresholding, denoising, and segmentation may enhance visual clarity or produce more precise outlines, none of these processes determine where a ratio is genuinely valid^9^ as a quantitative measure.

An optical solution, therefore, necessitates a set of contrasts with clearly defined evidentiary roles. Simultaneous label-free autofluorescence multi-harmonic (SLAM) microscopy captures NAD(P)H, FAD, and second and third-harmonic generation signals from a single broadband femtosecond excitation^10–14^, thereby providing metabolic intensity alongside nonlinear structural context. Quantitative phase imaging (QPI) assesses optical path length and produces detailed morphological data^15,16^ without inheriting the photon-support failure mode that compromises intensity ratios. Because phase does not depend on autofluorescence photon counts, it can independently interrogate cellular context without relying on the redox denominator. Finally, fluorescence lifetime imaging microscopy (FLIM) measures redox state via fluorophore binding^17–20^ and provides a lifetime-sensitive benchmark for validation. In this study, support is defined as the collective adequacy of local photon statistics and phase-derived cellular structure for reliable quantitative redox inference.

Here, we introduce FPhaS, a fixed-calibration phase–autofluorescence framework that seamlessly integrates quantitative phase imaging (QPI) with simultaneous label-free autofluorescence multi-harmonic microscopy (SLAM), using fluorescence lifetime imaging (FLIM) exclusively for validation. This methodology facilitates confidence-supported label-free redox imaging. Its fundamental design approach is intentionally restrictive; QPI is never utilized to replace or predict the redox signal. Instead, a one-time non-biological calibration maps the phase structure to the autofluorescence coordinate system, which remains fixed for all biological data, thereby ensuring that phase serves solely as a fluorescence-independent support criterion. The practical implementation is enabled by sharing the objective: the phase and autofluorescence fields occupy a single geometric space, allowing a single fixed calibration to align them with sub-pixel accuracy across all biological data. This setup permits phase to interrogate cellular context at the same pixel as the redox measurement. Based on this framework, FPhaS disaggregates cell detection, ratio admissibility, and confidence-supported inference into nested domains, benchmarks the resulting redox intensity estimates against FLIM-derived readouts confined to the mask, propagates support information into cell-level descriptors, and ultimately assesses whether the same declared support layer can restrict reduced-acquisition reconstruction.

## Results

### Fixed QPI-SLAM calibration links phase structure to metabolic readouts

FPhaS integrates its structural and metabolic measurements through a unified objective and a single sample coordinate system (Fig. 1a). The QPI arm provides a phase-derived structural map^15,21^. The SLAM arm records NAD(P)H, FAD, and second and third-harmonic channels^9-13^. FLIM operates on a nonlinear pathway using time-correlated single-photon counting^17–20^ and is intended to validate intensity-derived redox estimates; it is not incorporated into FPhaS as an input. This division of responsibilities constitutes the core of the framework: autofluorescence measures metabolism, phase supplies structural information independently, and lifetime data verifies the redox estimate retrospectively.

**Figure 1.**
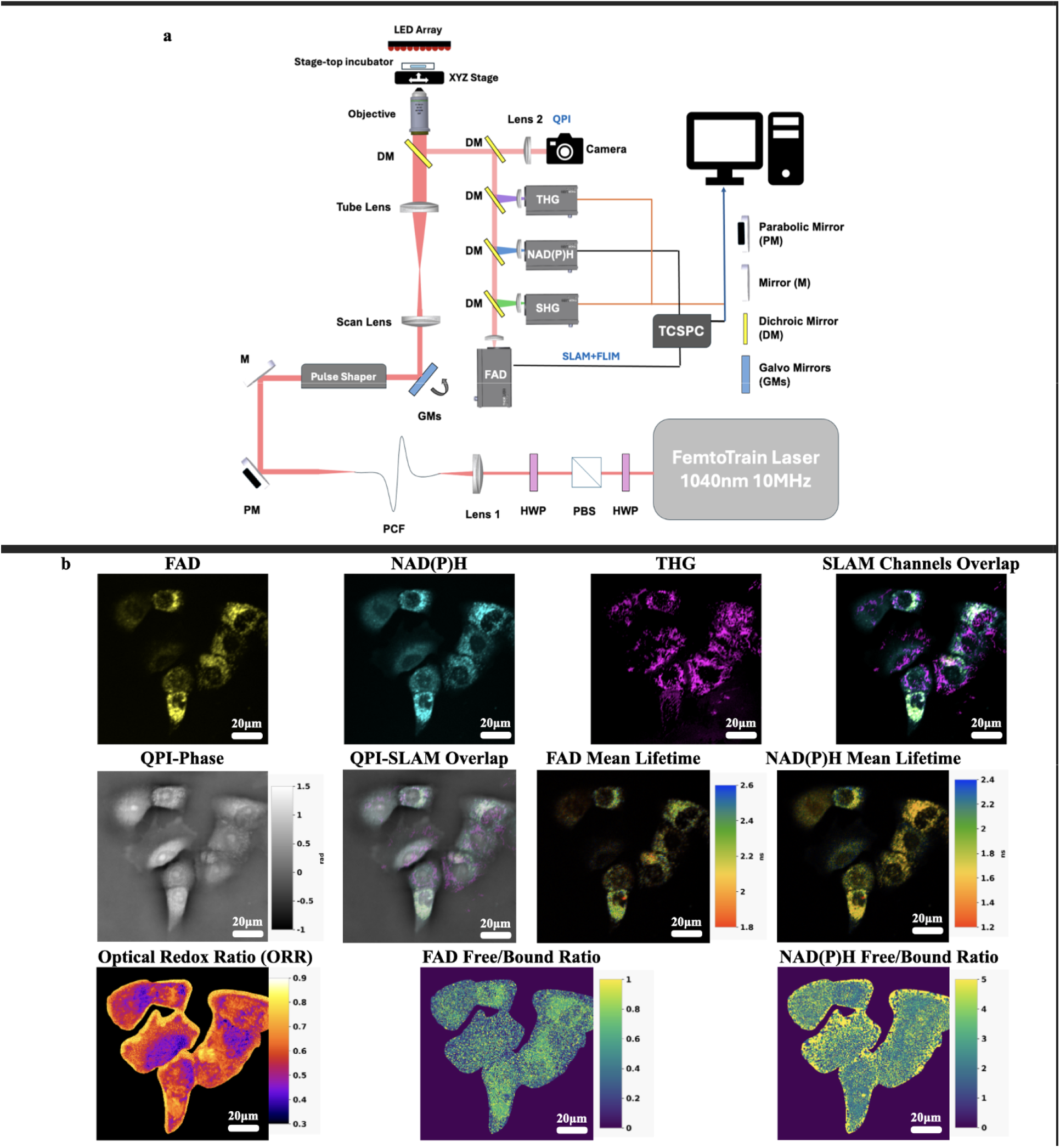
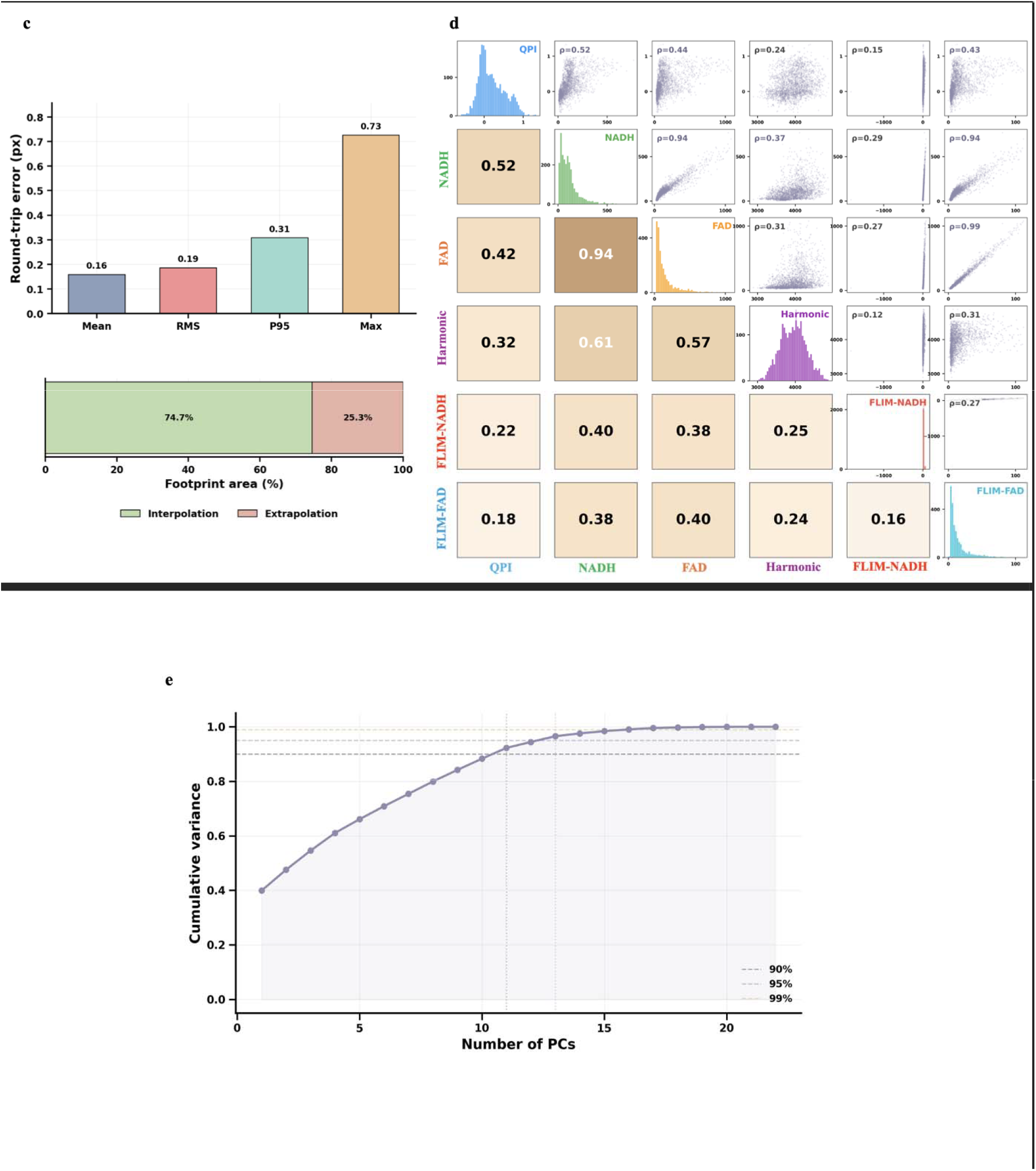
FPhaS establishes fixed phase–autofluorescence geometry and non-redundant structural information. a, Shared-objective acquisition and analysis schematic for co-registered QPI, SLAM and FLIM measurements. QPI provides phase-derived structure, SLAM provides NAD(P)H and FAD autofluorescence with harmonic-generation channels, and FLIM provides an orthogonal lifetime derived validation readout. b, USAF target calibration showing the SLAM side target image and the corresponding target mapped into QPI coordinates. c, Representative co-registered multimodal field acquired in the shared FPhaS geometry. The displayed channels include FAD autofluorescence, NAD(P)H autofluorescence, THG contrast, a SLAM composite image, FLIM derived FAD and NAD(P)H mean lifetime maps, the intensity optical redox ratio (ORR), FLIM derived FAD and NAD(P)H free/bound ratio maps and the matched QPI optical path length image(A-J). These maps illustrate local spatial correspondence among phase derived structure, autofluorescence contrast, and lifetime-derived redox context. FLIM derived maps are shown as validation context and were not used to construct the FPhaS confidence maps. d, Registration audit for the retained second-order polynomial transform; the upper panel summarizes footprint restricted residual error, and the lower panel shows interpolation and extrapolation coverage within the validated landmark footprint. e, Interchannel relationship summary across co-registered phase, autofluorescence and FLIM derived channels. f, Effective dimensionality analysis showing the number of principal components required to explain multimodal variance. The calibration used 61 distributed non-biological landmarks, and the selected transform was frozen for all biological fields.

Registration was incorporated into the methodological framework rather than being considered a superficial step. Paired USAF-target images were used in a single calibration process^22,23^, with the resulting transformation fixed for all subsequent biological fields(Fig. 1b). In contrast to assessments based on non-biological landmarks and geometry-based audits, various mappings—including affine, projective, second-order polynomial, and third-order polynomial were compared. The second-order polynomial transformation demonstrated the best compromise between registration accuracy and geometric stability (Supplementary Fig. S1).

FPhaS was first implemented in a shared multimodal imaging geometry in which SLAM autofluorescence, FLIM context, THG contrast, and QPI optical path length were acquired from the same field of view (Fig. 1c). The representative field illustrates the spatial correspondence among metabolic autofluorescence, phase-derived structure, and lifetime-derived redox context. The FLIM-derived maps are shown here for independent validation and were not used to construct the FPhaS confidence maps. To make this correspondence quantitative, QPI and SLAM were aligned using a fixed second-order polynomial transform selected by shifted-block cross-validation. The selected transform achieved a sub-pixel in-sample residual of 0.390 pixels and maintained a positive Jacobian determinant^23^ throughout the mapped region, indicating a locally orientation-preserving registration. The calibrated footprint covered approximately 74% of the image area within the convex hull of the control points (Fig. 1d). Source and destination-space inverse-consistency^23^ RMS errors, full transform-family model selection, cross-validation results, and geometry audits are provided in Supplementary Fig. S1.

This calibration establishes the operating geometry for all subsequent procedures. The structure derived from phase, autofluorescence intensity, and FLIM validation readouts is now unified within a single coordinate system, without the need for re-adjustment of the transformation relative to the biological images. Since both arms utilize the same objective, the geometric relationship between them remains sufficiently stable to permit a one-time calibration on a non-biological target, without subsequent refitting to biological images. This stability enables phase to serve as an external structural reference rather than a fluorescence-derived one, facilitating the comparison of structure and metabolism at identical locations.

### QPI provides complementary structural information to autofluorescence

A structural guide earns its place only if it adds information relevant to the specimen without simply echoing the fluorescence channels. We therefore examined marginal and partial relationships^24^ across the co-registered channel set. QPI showed a measurable marginal association with NAD(P)H, FAD, and the derived fluorescence channels; however, after de-confounding, those associations weakened, with the QPI-fluorescence pairs weakening most (Fig. 1e,f).

The dimensionality analysis yielded consistent results. For the four primary nonlinear channels—NAD(P)H, FAD, SHG, and THG—the multimodal feature set incorporated an additional seven FLIM-derived channels per fluorophore, as well as QPI-derived structural, gradient, and texture features. Eleven principal components accounted for 90% of the variance^25,26^ in the data (Fig. 1g), with the highest loadings indicating that the QPI variables form components distinct from those associated with NAD(P)H and FAD intensity (Supplementary Fig. S1).

FPhaS therefore considers QPI as a co-registered structural constraint rather than a substitute for NAD(P)H, FAD, or FLIM. Phase remains bound to the cellular specimen while concurrently conveying information that cannot be captured by autofluorescence intensity alone.

### FPhaS separates cell presence from inferred metabolic support

After establishing the shared FPhaS geometry and fixed QPI–SLAM registration. We next examined how structural and autofluorescence support differed within registered biological fields. QPI provided fluorescence-independent structural evidence for cellular material, whereas the NAD(P)H and FAD intensity channels determined where the optical redox ratio could be formed with adequate autofluorescence support. These support sources were spatially correlated but not interchangeable: phase-defined cellular structure did not always provide reliable autofluorescence-ratio support, and fluorescence-positive regions did not automatically satisfy the structural-support criterion. FPhaS therefore defined nested support domains in which pixels were classified as QPI-supported, fluorescence-supported, or confidence-supported. A pixel is deemed supported by QPI when its phase-based structural reliability score surpasses a predetermined threshold. It is considered supported by fluorescence when both its total autofluorescence denominator and the propagated redox ratio uncertainty^27,28^ satisfy predefined support criteria. A pixel is classified as confidence-supported only when both conditions are simultaneously met. This hierarchical nesting of domains segregates three critical decisions—namely, the presence of cellular material, the feasibility of ratio evaluation, and the adequacy of joint support for pooling—processes that are typically conflated in a single mask (Fig. 2).

**Figure 2.**
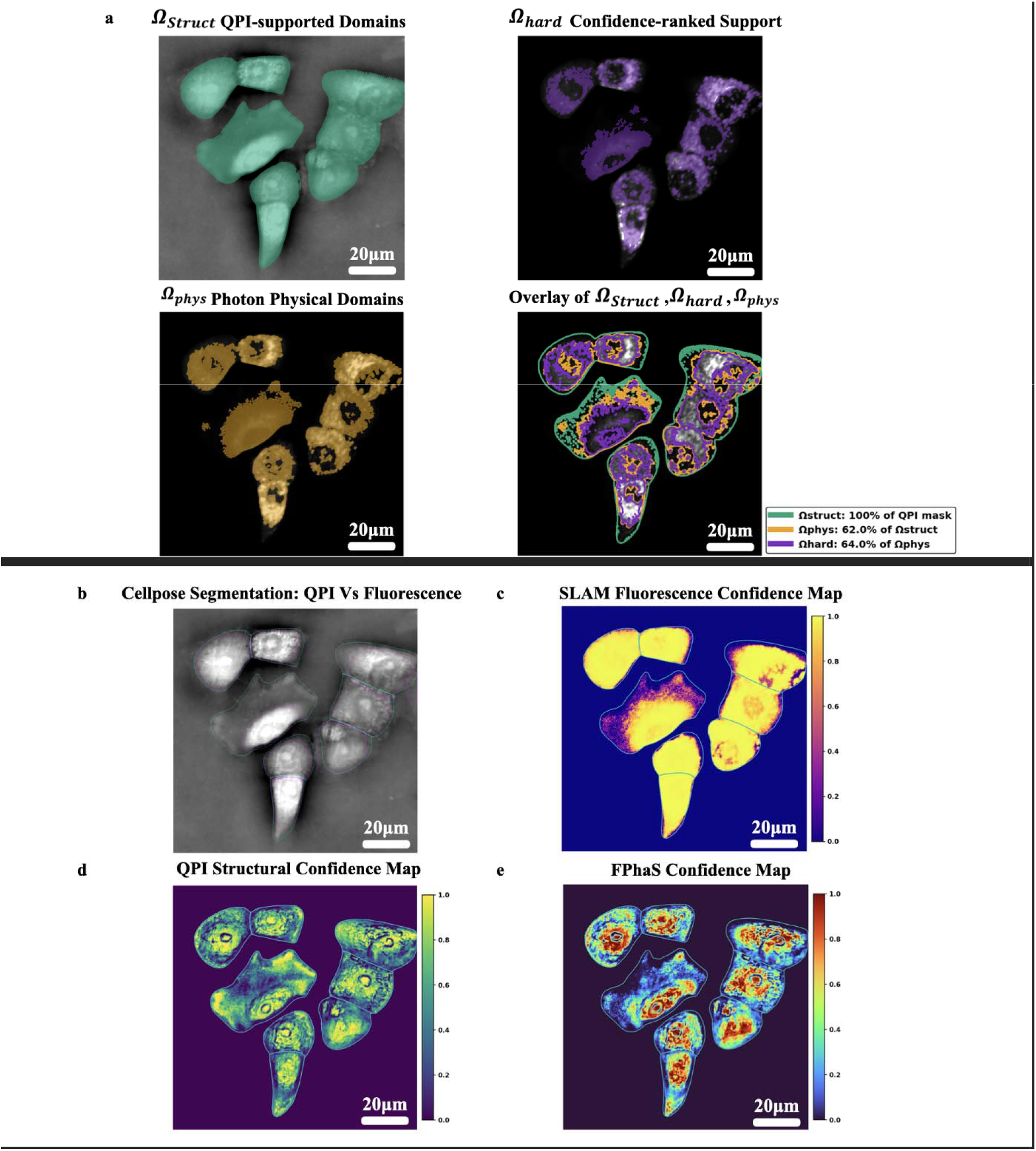
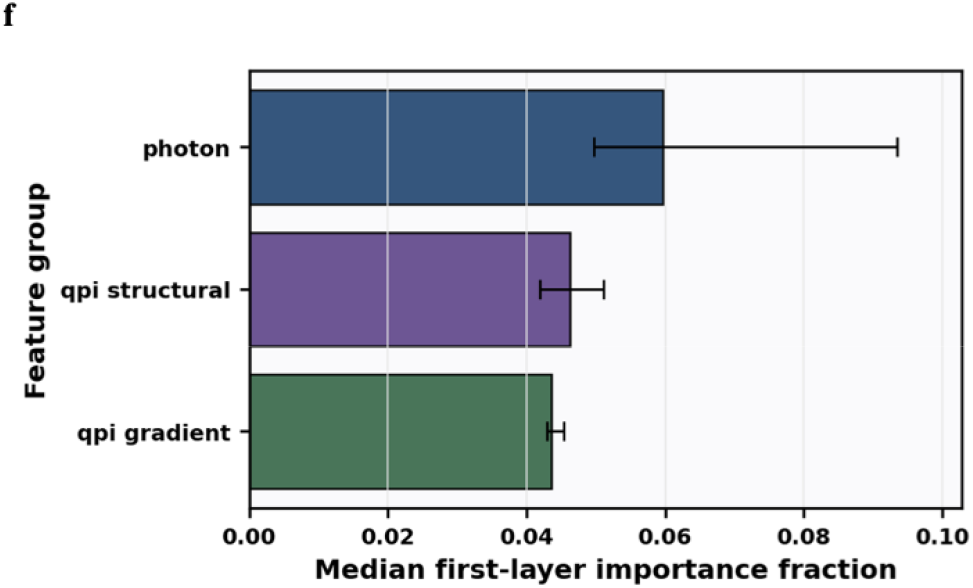
FPhaS links co-registered phase–autofluorescence measurements to confidence-supported redox inference. a, b, Registered support domain construction in the same field. The internal panels show QPI supported cellular structure, SLAM derived fluorescence support, raw intensity ORR and the overlay of structural, physical and hard operating domains(A-D). These maps separate cell presence, ratio admissibility and confidence supported operating support. c, QPI phase image used to define structural cellular support. d, SLAM denominator-support image, defined as NAD(P)H + FAD. e, QPI derived structural confidence map. f, Autofluorescence support confidence map derived from denominator support and local photon support. g, Primary FPhaS confidence map combining phase-derived structural support with autofluorescence support inside the declared operating domain. h, Feature group importance for photon, QPI structural and QPI gradient features. The confidence field ranks pixels within declared support domains rather than converting unsupported pixels into quantitative evidence.

The confidence metric integrates fluorescence-based support with structural support derived from QPI, with each component addressing distinct failure modes. The fluorescence component assesses the denominator support^27–29^ for the redox-ratio estimator; a pixel may appear to have a valid ratio but provide minimal quantitative data if its autofluorescence denominator is weak. The QPI component assesses an alternative aspect, indicative of phase-derived cellular structure and local context. A pixel exhibiting high fluorescence intensity may lack structural clarity, whereas a pixel that appears structurally reliable may suffer from photon starvation. Consequently, the combined score evaluates two independent dimensions of reliability, rather than condensing all information into a single brightness metric.

Feature-group analysis substantiates a dual-source reliability perspective. The median per-feature permutation importance (the decrease in confidence-model performance when that feature is randomly permuted) was 0.0556 for photon features, 0.0544 for QPI structural features and 0.0515 for QPI-gradient features (Fig. 2g). These values do not represent cumulative totals for the groups; rather, importance is distributed across numerous features within each group. The proximity of these median values aligns with a model in which fluorescence support and phase-derived structural information collaboratively contribute to the confidence field, while remaining distinct, non-interchangeable sources of evidence.

The same distinction was observed at the image level. Masks derived from QPI and SLAM overlapped but did not fully coincide, with QPI masks indicating structural support and SLAM-derived masks indicating fluorescence contrast. Raw ORR maps contained regions where the total-fluorescence denominator support was weak, even when the displayed ratio appeared plausible, and FPhaS specifically identified these regions as low-confidence for primary quantification.

This methodology ensures a clear delineation between exclusion and weighting. The denominator-support rule initially determines the locations where the ratio estimator possesses sufficient measurement support; subsequently, confidence weighting ranks the pixels within that set. The safeguard integral to the design is straightforward: unsupported pixels are never elevated to the status of quantitative evidence post-acquisition. They are excluded from primary inference and are not salvaged through denoising, interpolation, or display smoothing.

### Mask-locked FLIM validation defines the primary performance metric

Validation in this context is more nuanced than it initially appears. Comparing intensity and lifetime readouts across various masks primarily assesses segmentation accuracy. This potential pitfall was circumvented by employing FLIM-derived redox information as an external benchmark^17–20^ for the support model within cell regions defined by masks (see Fig. 3). The FPhaS confidence map was constructed solely from the QPI and autofluorescence channels. Only subsequently did we evaluate its concordance with FLIM-derived redox readouts in the identical cellular regions, ensuring that the performance measure reflected reliability ranking within a shared support domain rather than differences in pixel inclusion.

**Figure 3.**
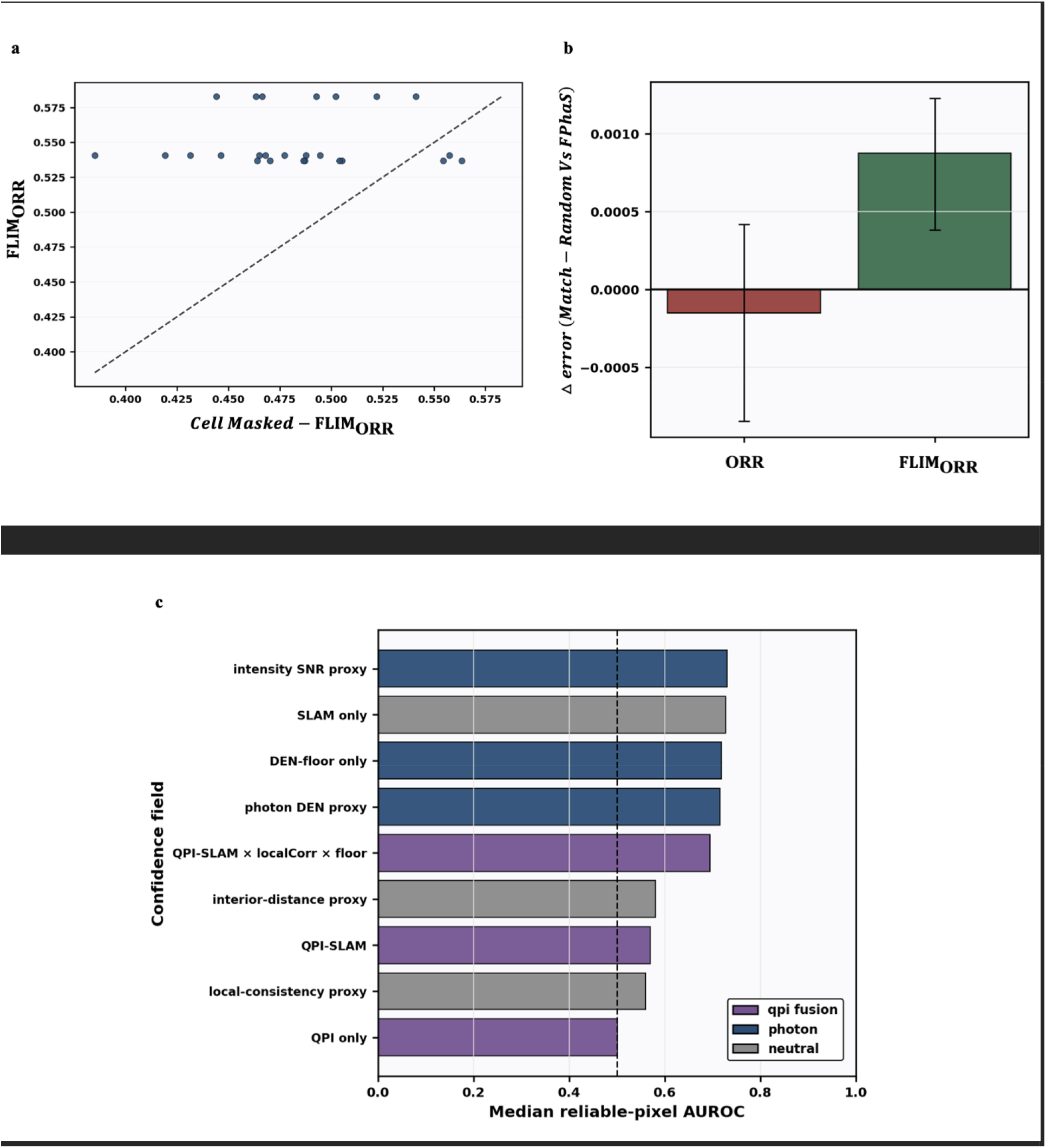
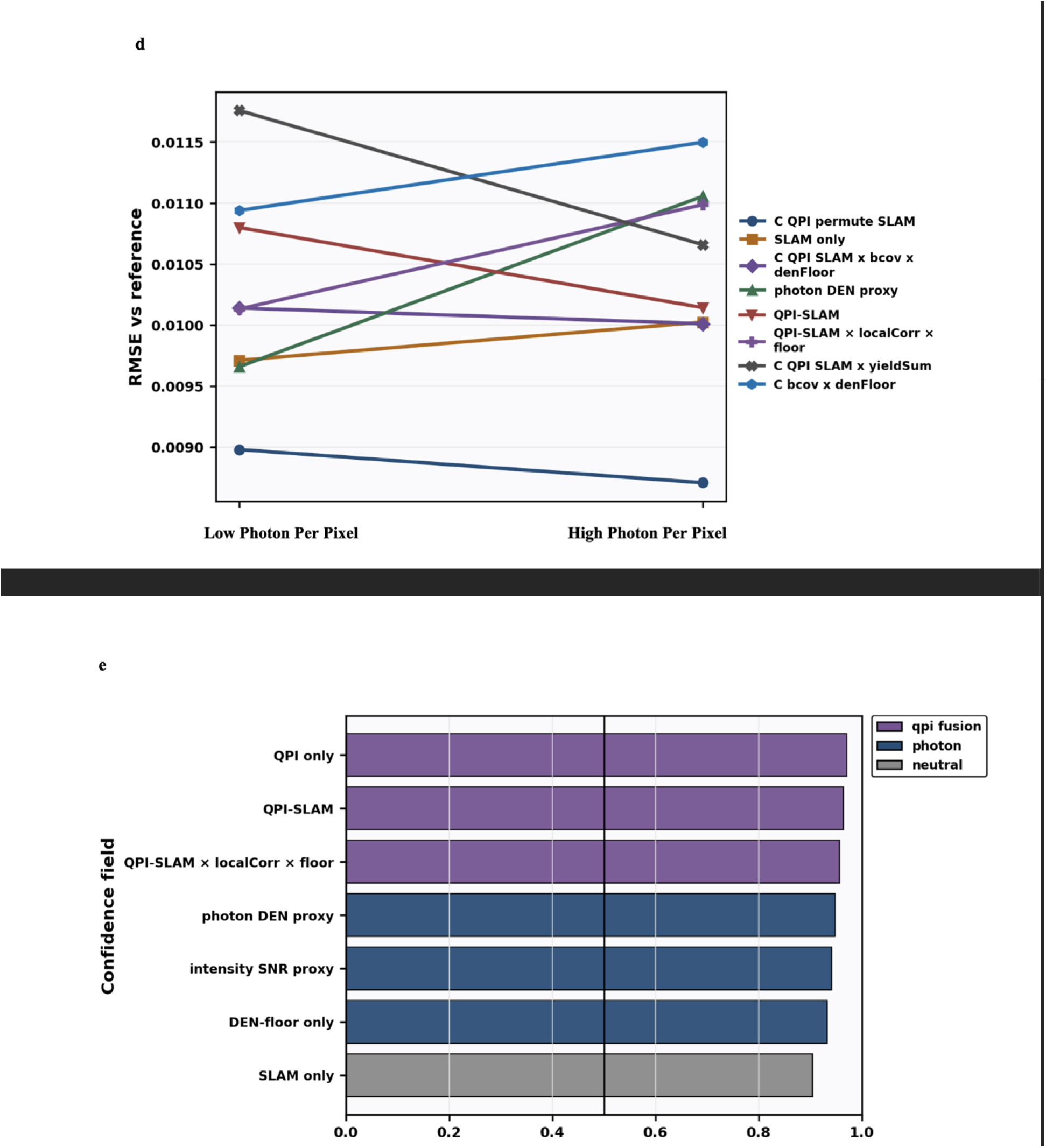
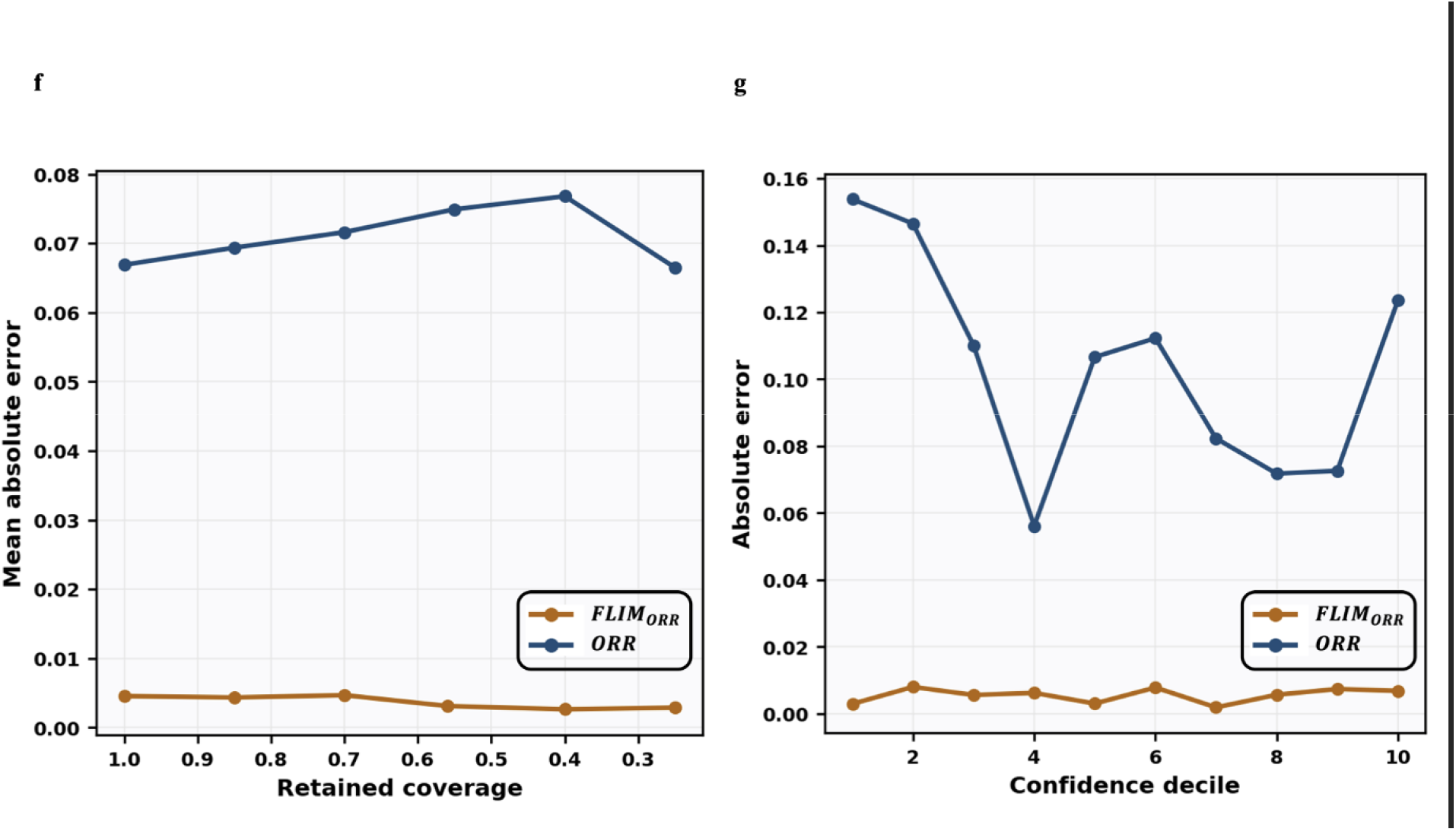
Mask-locked FLIM-derived redox validation defines the FPhaS performance boundary. a, Mask lock audit comparing FLIM derived redox estimates computed inside QPI supported cellular masks with FLIM-derived redox estimates computed from the full image context. The dashed line denotes equality. b, Denominator matched random-control analysis for intensity redox and FLIM derived endpoints. Values show Δ error, defined as matched-random error minus FPhaS error; positive values indicate lower error for FPhaS. c, Pixel confidence ranking for selective quantification. d, Photon support robustness under high and low photon per pixel conditions. e, Confidence field comparison across photon only, QPI only, SLAM only and fused selectors. f, Risk coverage behavior after confidence based pixel selection. g, Confidence monotonicity analysis. FLIM derived redox information was used as an orthogonal validation readout to test whether FPhaS confidence maps improve intensity-based redox agreement within a shared cellular support domain, not across mismatched masks.

A mask-lock audit confirmed that the intensity and FLIM-derived redox estimates were computed on the same QPI-supported pixels within each analyzed cell. The validation set comprised 10 fields of view across 5 samples, with both high and low-photon analyses included to stress-test robustness to photon support. Applying the mask shifted the reference calculation relative to the full-image context (median absolute difference of 0.0667; Fig. 3a), confirming that the endpoint under validation was genuinely cell-localized rather than a field-level average in disguise.

Denominator-matched control^30^ groups enable the differentiation between confidence-based selection and denominator selection. Regarding the comprehensive-field intensity redox endpoint, the initial confidence map did not outperform the denominator-matched control groups (mean random-minus-primary error difference: −1.5 × 10□□; 95% Confidence Interval crossing zero). This negative outcome is significant: FPhaS is not a universal enhancer across all redox maps presented. Conversely, the mask-locked endpoint yielded a different result. In this case, the primary confidence map demonstrated a smaller absolute error compared to the denominator-matched controls (mean random-minus-primary error difference: 8.75 × 10□□; 95% Confidence Interval, 3.82 × 10□□ to 1.227 × 10□^3^; see Fig. 3b). In summary, the observed gain is specific to the endpoint and is associated with the designated cellular support domain.

Pixel-level analyses aligned with this reading. Confidence ranking, spatial stability, and risk-coverage behavior together provided a reliability ordering among supported pixels (Fig. 3c-g), though we did not treat the confidence-decile trend as a separate biological readout. On this evidence, the validated gain lies in support-aware quantification, not in image beautification or in any post-acquisition recovery of unsupported measurements.

High and low-photon analyses adhere to identical principles under conditions of diminished photon support. As the number of photons per pixel decreases, the confidence model consistently distinguishes between pixels defined solely by ratios and those supported by independent photon and structural evidence. Consequently, the practical output of FPhaS is not intended to produce a more comprehensive redox image; rather, it defines a specified operating domain for redox quantification.

### MCI, MOI, and MRI propagate support information to cell-level aggregation

FPhaS next extends support-aware inference from pixels to entire cells (Fig. 4). Aggregation is precisely where support information tends to diminish: two cells with nearly identical mean redox ratios might still differ in the localization of the supported signal, its alignment with phase-derived structures^15,16^, and the extent to which effective sample support^31^ persists after weighting. The descriptor analysis examines whether these differences can be preserved once the data are aggregated at the cellular level.

**Figure 4.**
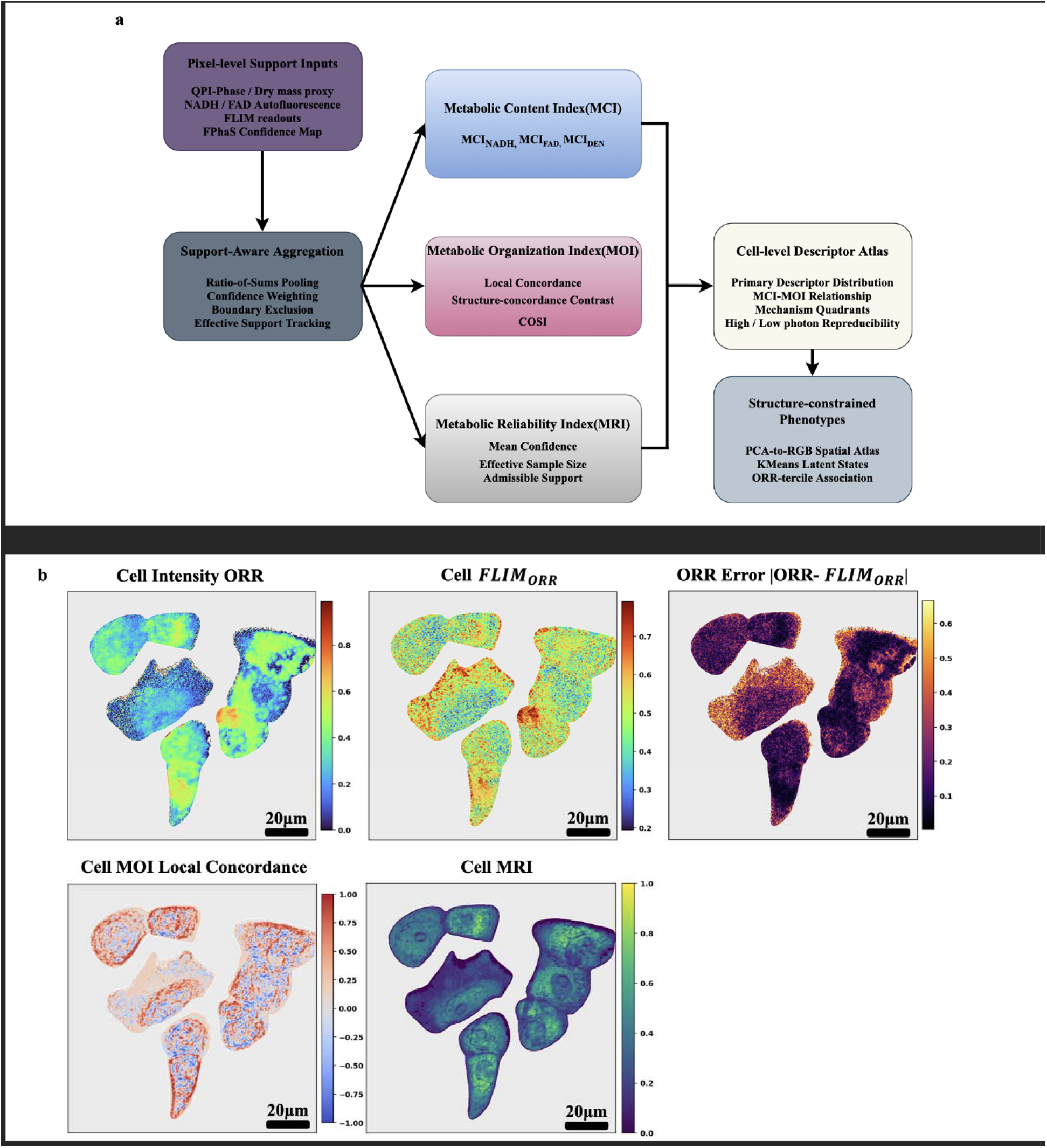
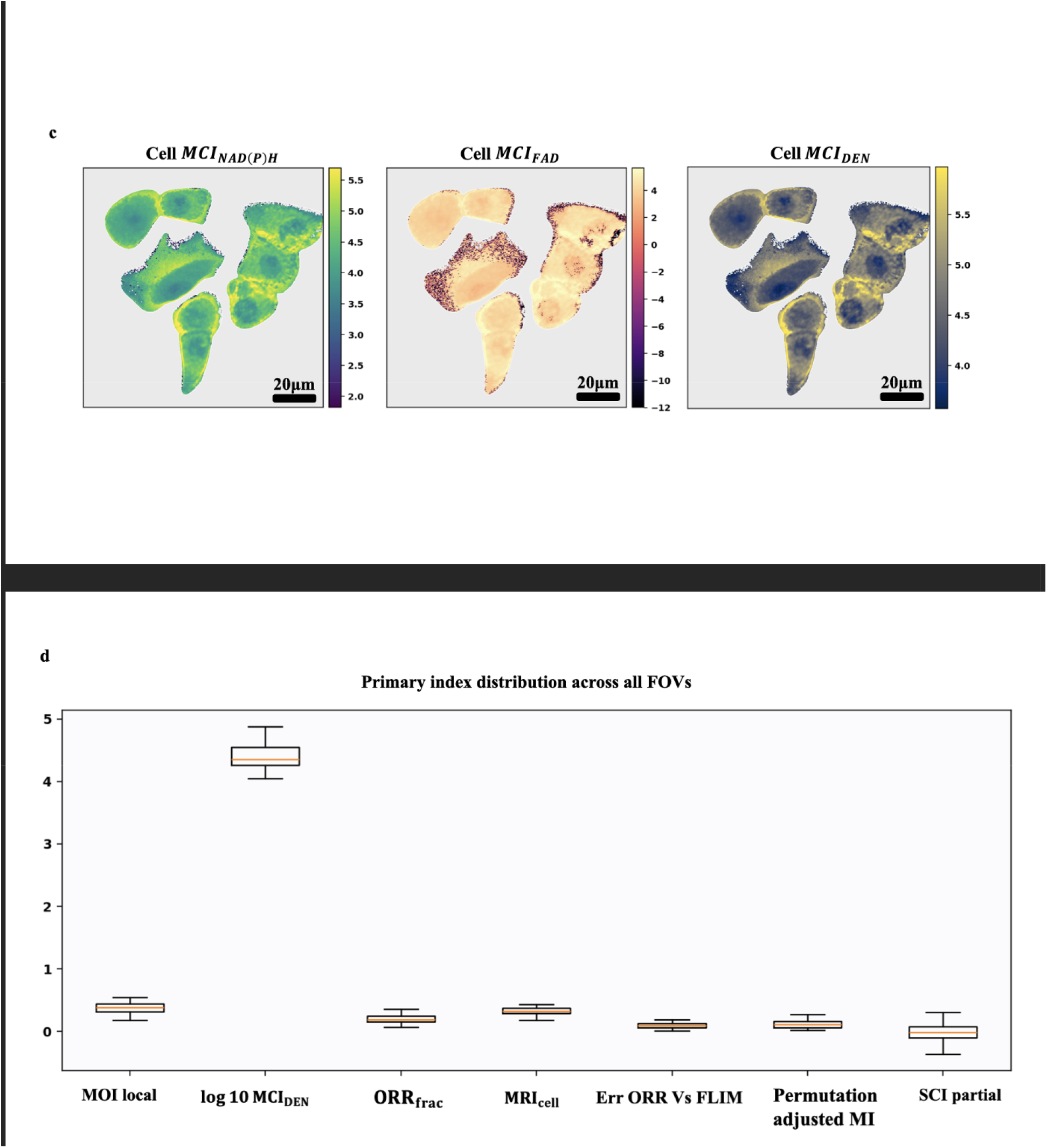
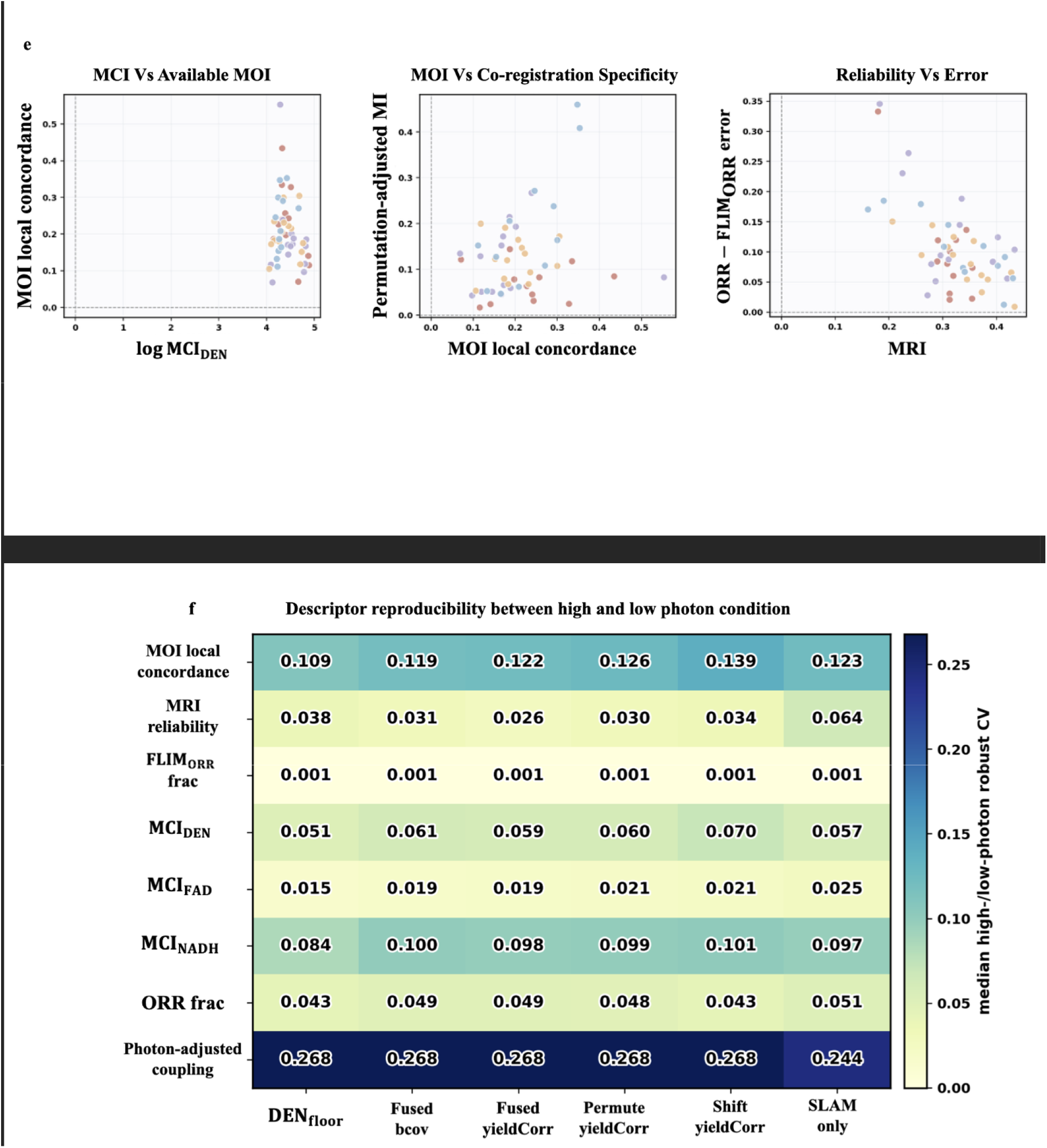
MCI, MOI and MRI preserve support information during cell-level aggregation. a, Descriptor definition schematic. Pixel level QPI–SLAM measurements are restricted to the hard operating domain and aggregated with FPhaS confidence weights. MCI summarizes confidence-weighted metabolic content over QPI derived structural support. MOI summarizes metabolic–structural organization, including local concordance, structure-conditioned contrast and permutation-adjusted specificity. MRI summarizes confidence, admissible support and effective sample support within each cell. b, Representative cell-level descriptor atlas. Internal panels show cell level maps of support-aware redox and descriptor quantities derived from the same field. FLIM derived maps, where shown, are included only as external diagnostic references and were not used to construct MCI, MOI or MRI (A-F). c, Channel specific MCI maps showing mass-normalized metabolic yield for NAD(P)H, FAD and denominator derived channels(A-C). MCI values are relative fluorescence per structural support indices and are reported in arbitrary units. d, Primary descriptor distributions across analyzed cells. MOI and MRI are dimensionless descriptors; mutual information based MOI terms are reported in bits when computed with log2. e, Support aware descriptor relationships. The internal panels show the MCI–MOI relationship, MOI local concordance versus permutation adjusted mutual information, and MRI versus ORR–FLIM error. Permutation adjusted mutual information is computed as observed QPI-mass–ORR mutual information minus the within-cell permuted null. FLIM derived ORR is used here only as an external diagnostic reference. f, Descriptor reproducibility between high and low photon conditions. g, Joint descriptor embeddings and loading summaries. The internal panels show PCA organization, UMAP organization and descriptor loading structure, illustrating how MCI, MOI and MRI contribute to support aware cell level variation. The analysis included 10 fields of view across 5 samples.

We established three descriptor categories. The Metabolic Content Index (MCI) encapsulates the permissible metabolic burden in relation to the structural support derived from QPI. The Metabolic Organization Index (MOI) evaluates the degree to which the redox signal is tightly coupled to phase-derived structures. The Metabolic Reliability Index (MRI) summarizes the confidence distribution and the effective sample support^31^ within each cell. None of these indices aims to define a specific cell state; rather, they serve as support-preserving summaries that coexist with and complement the overall cell redox value.

The descriptor analysis utilized the same dataset (10 fields of view, 5 samples). MCI, MOI, and MRI were all computed within the primary index family. The median MCI was 22,573.5, the median MOI local correlation was 0.188, and the median MRI was 0.319 (Fig. 4c-e). MCI values are relative fluorescence-per-structural-support indices and are reported in arbitrary units because the current implementation uses background-corrected autofluorescence intensity and a QPI-derived dry-mass proxy rather than absolutely calibrated fluorophore concentration and dry mass. MOI and MRI are dimensionless descriptors; mutual-information-based MOI terms are reported in bits when computed with log2. Collectively, these values uphold the distinctness of metabolic burden, structural organization, and measurement reliability as cell-level quantities.

We then checked descriptor stability between high and low-photon conditions. Variability remained consistently low, with a median coefficient of variation of 0.0537 (Fig. 4f). Confidence-weighted descriptors therefore remained measurable despite the decrease in photon support, indicating that support-aware aggregation effectively preserves valuable cell-level structure without assuming that every pixel within a segmentation mask contributes equally.

In the joint descriptor space, cells are categorized by burden, organization, and reliability (Fig. 4g). Cells with similar mean redox ratios do not necessarily cluster within the same descriptor neighborhood, a variation that reflects genuine differences in support and spatial arrangement. Consequently, MCI, MOI, and MRI are not redundant paraphrases of a whole-cell average. They preserve information regarding the distribution of the metabolic signal across the structurally supported regions of each cell.

### Confidence-weighted descriptors organize reproducible, structure-constrained cell variation

Next, we asked whether the confidence-weighted descriptor panel could systematically organize cell-level variation across different fields (Fig. 5). Principal component analysis was employed to encapsulate the descriptor space directly, without the preliminary step of passing raw pixel maps to the clustering procedure. The primary axes captured variations in metabolic burden, metabolic-structural organization, and reliability structure (Fig. 5a,b).

**Figure 5.**
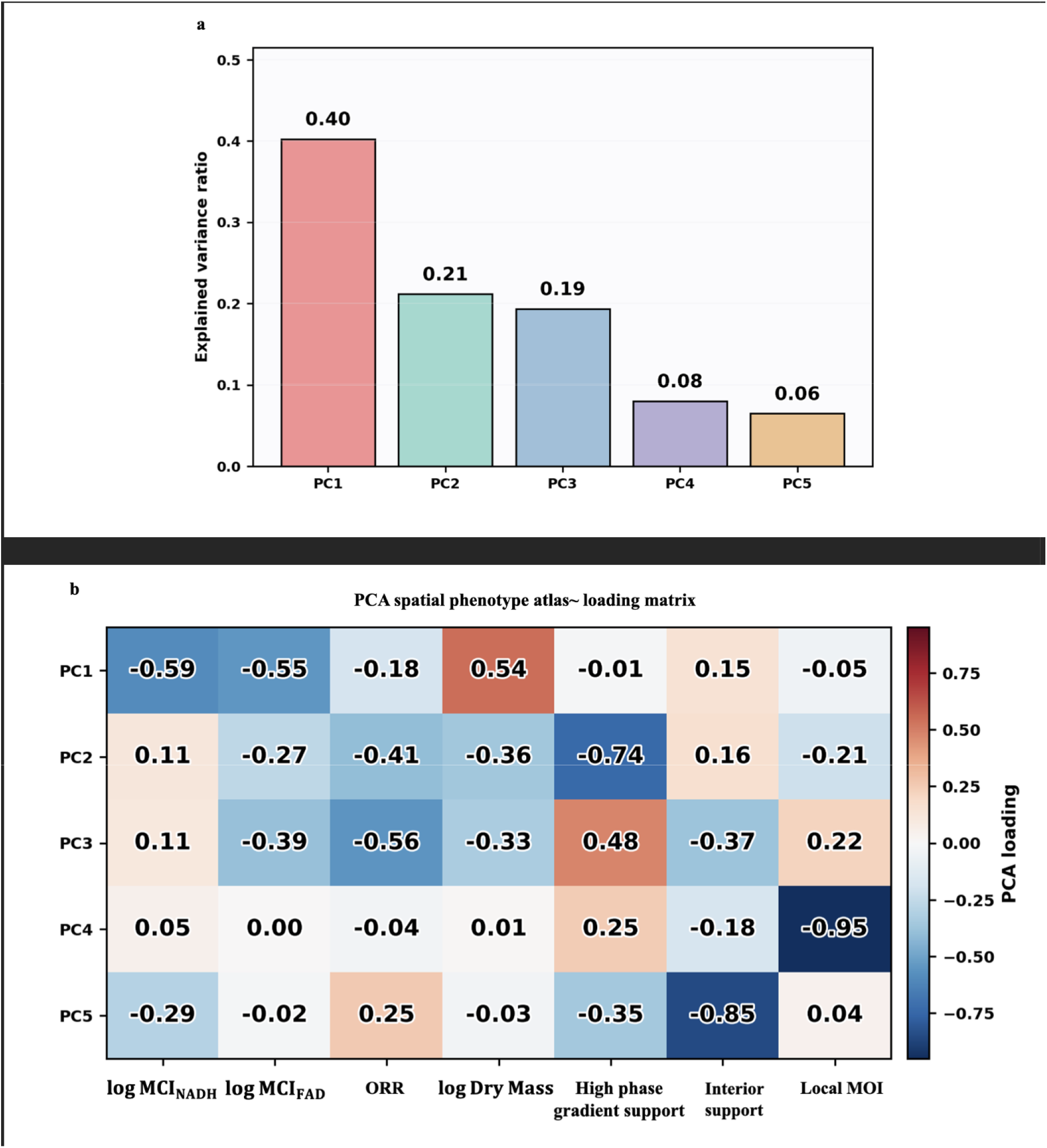
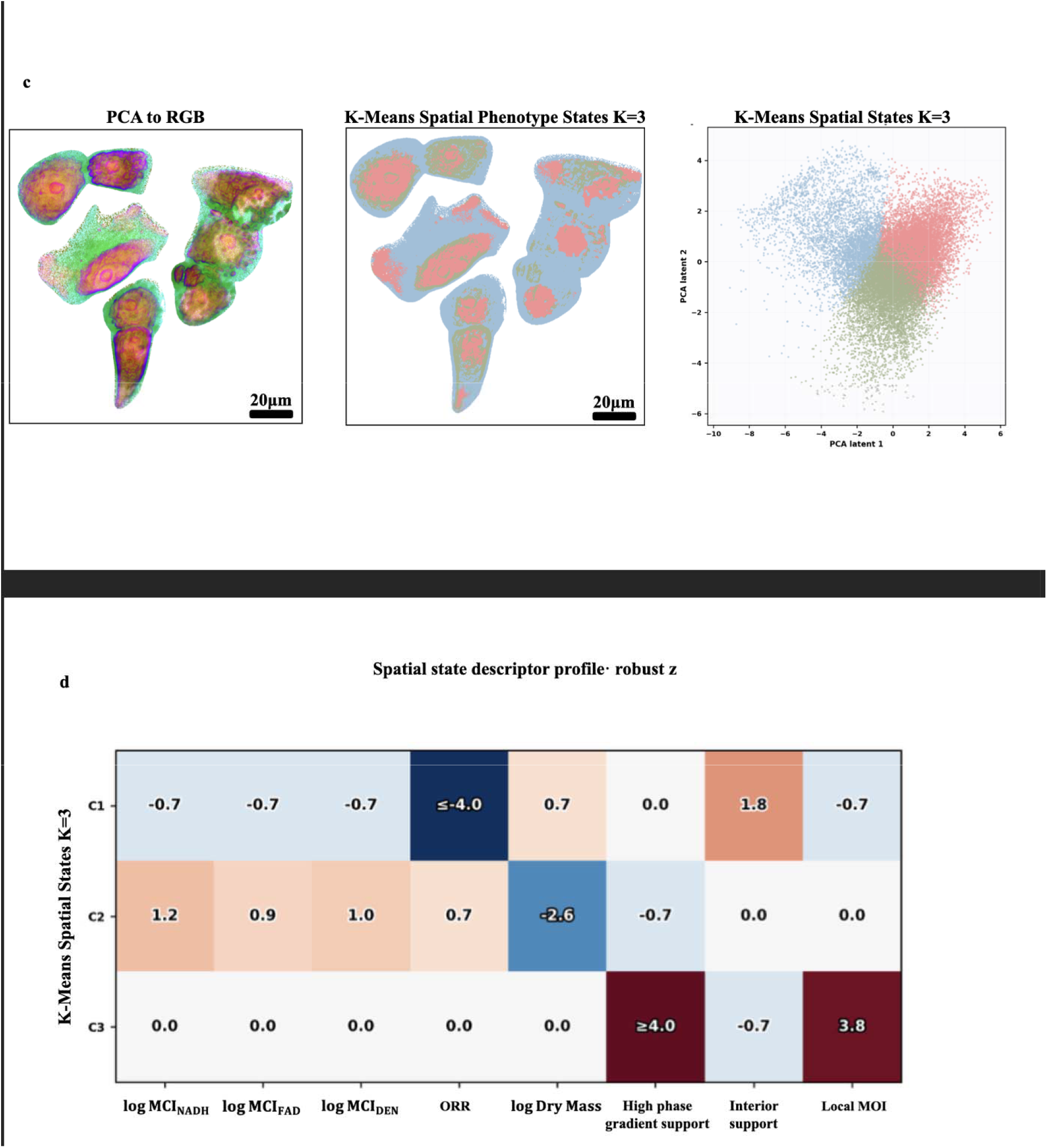
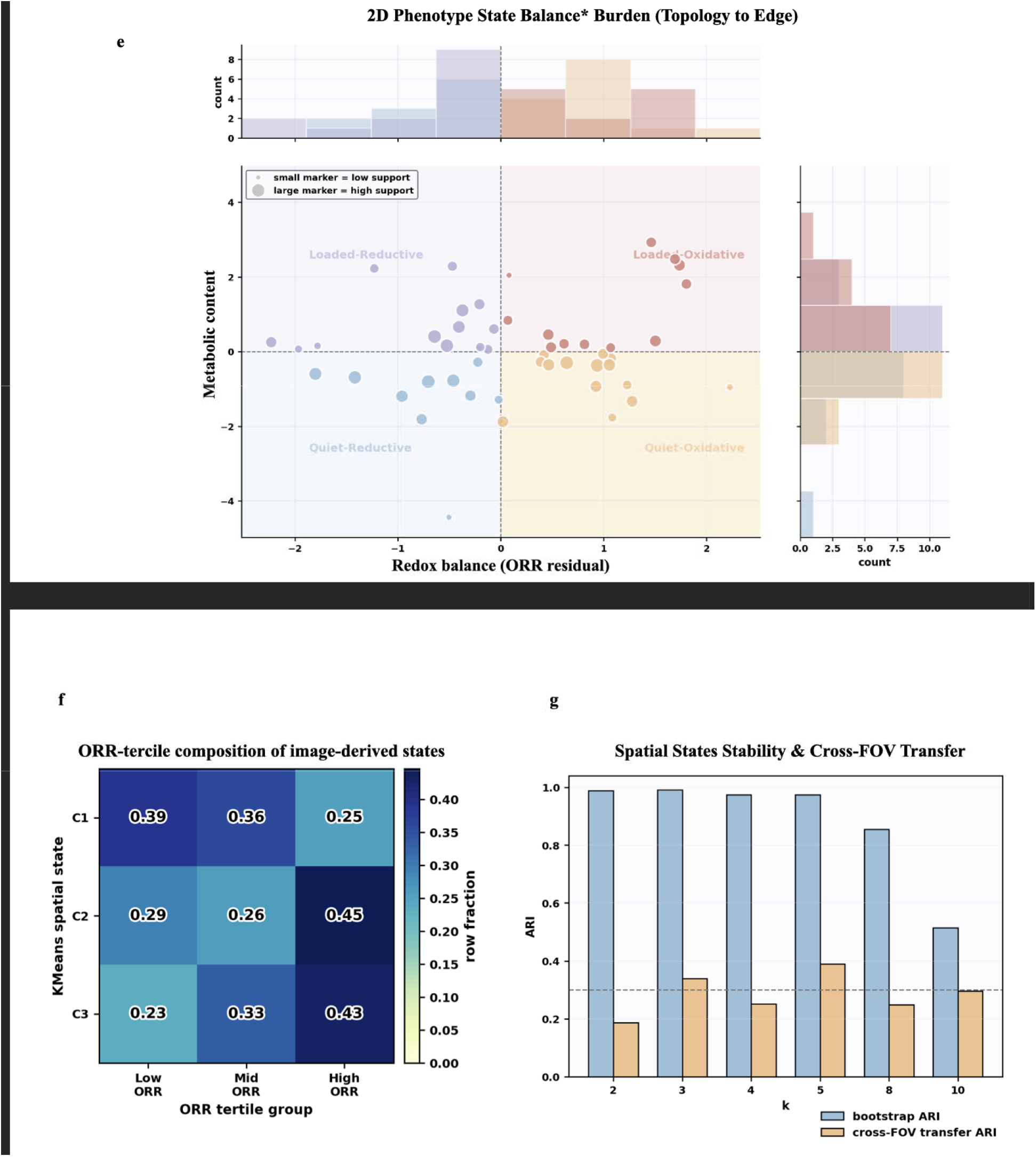
Confidence-weighted descriptors organize reproducible, structure-constrained cell variation. a, Principal component variance explained by the confidence weighted descriptor panel. b, Principal component loading heatmap showing how MCI derived metabolic content, ORR, QPI derived dry mass, phase gradient support, interior support and local MOI contribute to the phenotype axes. c, Descriptor derived spatial organization. Internal panels show the PCA-to-RGB spatial map, the corresponding image derived state map and the latent space distribution used to visualize three state organization. d, State level descriptor profile showing how MCI derived content, ORR, QPI structural support features and local MOI vary across image derived states. e, Descriptor state composition in metabolic content and redox balance space. Cells are plotted by residualized ORR and *MCI*_*DEN*_ axes. Positive ORR residuals indicate oxidative shifted cells, whereas negative residuals indicate reductive shifted cells. Positive *MCI*_*DEN*_ residuals indicate higher metabolic content, whereas negative values indicate lower metabolic content. Marker size indicates effective support, and marginal histograms show the distribution across descriptor quadrants. f, Association between descriptor states and ORR terciles. g, Descriptor state stability, cross field transfer and state coverage summary. The state map is interpreted as an image derived organization of the sampled fields, not as a fixed biological taxonomy.

Mapping the descriptor principal components^25,26^ to RGB revealed spatially organized variation within the sampled fields (Fig. 5c). Three-state clustering in the descriptor latent space then produced an image-derived organization map (Fig. 5d). Each state profile retained contributions from MCI, MOI, and MRI rather than collapsing onto a single mean-intensity axis (Fig. 5e).

The three-state representation persisted after resampling, with a mean bootstrap-adjusted Rand index^32,33^ of 0.991. Cross-field transfer demonstrated reduced strength but remained above chance level, with a mean adjusted Rand index^33^ of 0.339. The minimum state coverage was recorded at 0.174. The association with redox state persisted through permutation testing, with a modest effect size of Cramér’s V = 0.147 (Fig. 5f-h).

We interpret these results as evidence that the descriptor map serves as a reproducible summary of structure-constrained cell variation across the sampled fields. They do not constitute a fixed biological taxonomy. Cross-field transfer and redox-state association were modest, and we did not include direct QPI-disruption controls. For that reason, we present the state map as an illustration of support-preserving aggregation, rather than as definitive proof of discrete metabolic cell types.

### FPhaS reconstruction preserves fluorescence readouts under reduced acquisition burden

The reconstruction branch asks a final question: can the declared support layer also serve to constrain the analysis of reduced-acquisition data? (Fig. 6) In this context, the low photon per pixel’s fluorescence image provided the measured metabolic signal, whereas the high photon per pixel’s fluorescence image served only as a benchmark reference. The co-registered QPI channel served as a registration anchor and an edge-aware regularizer^34^ within the support layer, not as an independent source of metabolic contrast. In essence, the reconstruction was required to preserve the measured fluorescence readouts within a specified domain, rather than to infer metabolism from phase alone.

**Figure 6.**
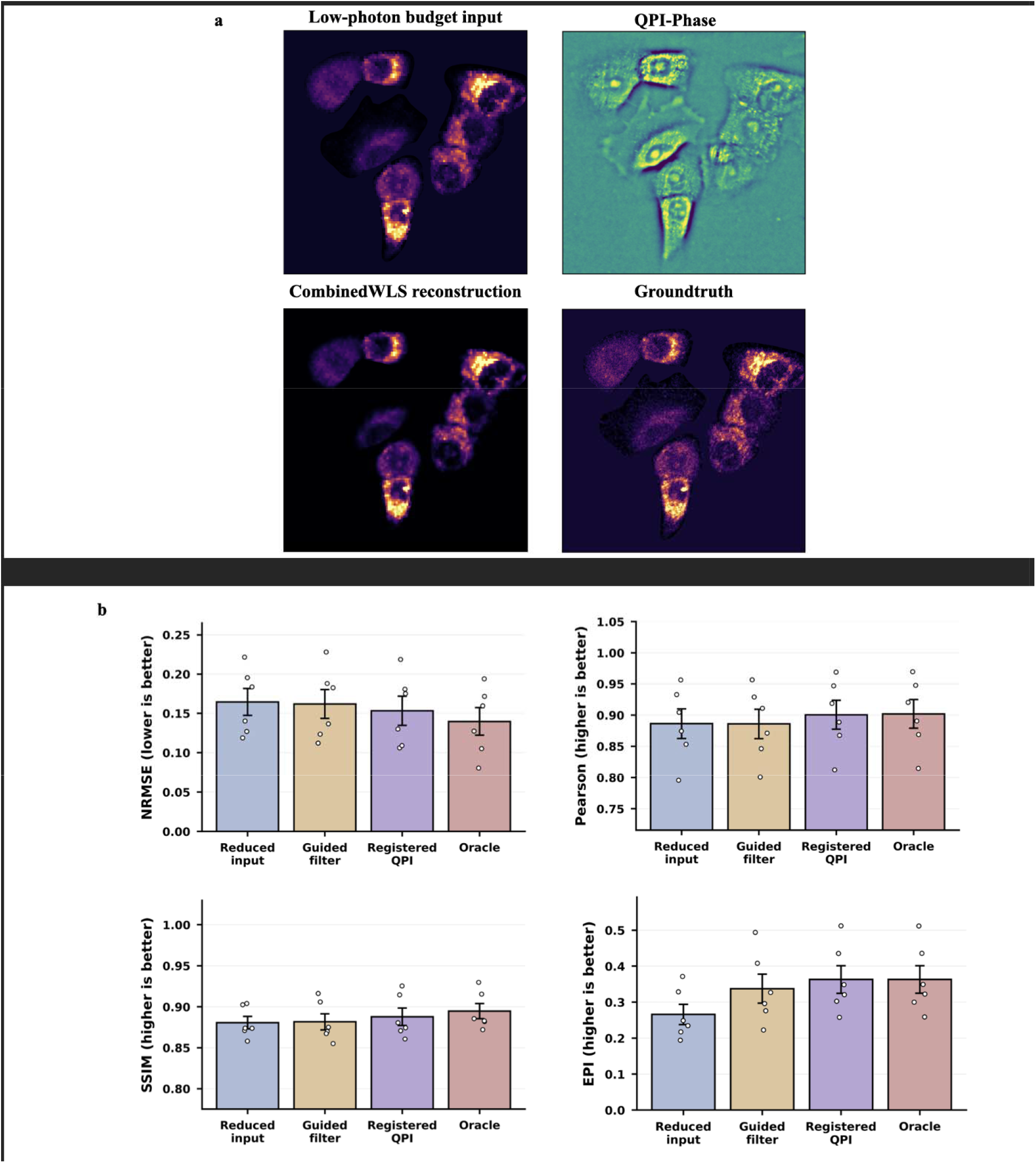
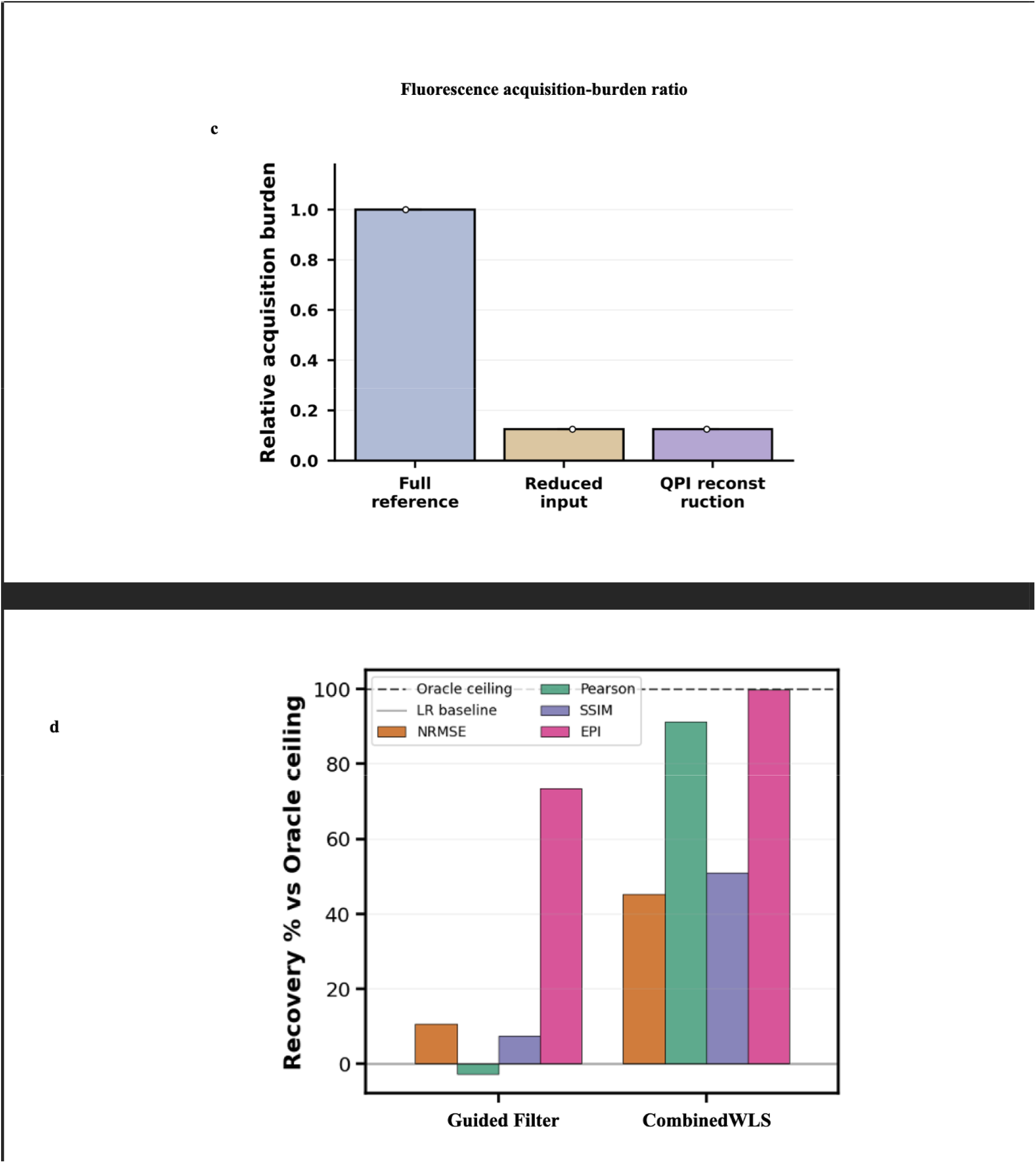
FPhaS reconstruction preserves fluorescence readouts under reduced acquisition burden. a, Representative low photon condition acquisition reconstruction. Internal panels show the low photon fluorescence input, co-registered QPI guide, CombinedWLS reconstruction and high photon condition acquisition reference. The low photon fluorescence image provides the measured signal, whereas QPI provides spatial support for the reconstruction regularizer. b, Support layer reconstruction benchmark. Internal panels show NRMSE, Pearson correlation, SSIM and edge preservation index for the low photon input, guided filtering, CombinedWLS and oracle upper bound. Lower values indicate better performance for NRMSE, whereas higher values indicate better performance for Pearson correlation, SSIM and edge preservation. Interpolation, Richardson–Lucy and calibration-ablation controls are reported in the Supplementary Information. c, Acquisition-burden analysis showing the eightfold low photon condition acquired fluorescence content per channel relative to the high photon acquisition reference under the tested protocol. d, Per-file paired improvement from low photon condition input to CombinedWLS reconstruction for NRMSE and Pearson correlation. e, Oracle gap recovery showing the fraction of possible improvement recovered by CombinedWLS across NRMSE, Pearson correlation, SSIM and edge preservation metrics. Registered QPI guided filter use and specificity controls, frequency resolved analyses and line profile diagnostics are provided in Supplementary Figs. S8 and S9.

Within the support layer, CombinedWLS improved every metric relative to the low photon per pixel’s image input, with the most significant enhancements observed in structural metrics (Fig. 6b). It increased the edge-preservation index by over one-third (0.266 to 0.363) and nearly halved the block artifacts (1.024 to 0.558), while reducing the support-layer reconstruction error by 7% (NRMSE 0.1645 to 0.1532). The improvements in the bounded agreement metrics were smaller in absolute terms (Pearson 0.886 to 0.901, SSIM 0.881 to 0.888), which is attributable to limited remaining headroom, as clarified by the oracle-gap analysis presented below. Control experiments using guided filtering^35^ configurations (no-QPI, shifted-QPI, and registered-QPI) and the baseline methods of interpolation and Richardson-Lucy^36,37^ in the Supplementary Information confirm that QPI acts as an edge-aware regularizer^34^ rather than a source of metabolic contrast.

Under the tested protocol, the reduced-acquisition condition had eightfold less acquired fluorescence per channel than the full-acquisition reference (Fig. 6c). After CombinedWLS reconstruction, the support-layer fluorescence metrics approached those of the reference. We therefore treat this branch as a reduced-information reconstruction benchmark, not as a direct measurement of wall-clock time, phototoxicity or viability unless those quantities are logged prospectively.

The per-file paired analysis showed that the improvement was not attributable to a single flattering example (Fig. 6d). Across individual files, CombinedWLS demonstrated enhancement over the low photon per pixel input condition for the primary accuracy and agreement metrics. Paired consistency is particularly important in this context, as, under conditions of reduced information, aggregate averages alone may obscure genuine file-to-file variability.

An oracle-gap analysis quantified the remaining progress (Fig. 6e). CombinedWLS achieved 45.3% of the available NRMSE reduction, 91.4% of the attainable Pearson correlation enhancement, 51.0% of the available SSIM improvement, and 99.9% of the available edge preservation enhancement relative to the oracle benchmark. The most substantial retained performance was in correlation, structural similarity, and edge preservation, whereas residual amplitude calibration persists as a limitation.

## Discussion

FPhaS addresses a support problem that typically remains concealed, bringing it into the open. Although a label-free redox ratio is straightforward to display, its numerical value alone serves as a weak standard for quantitative evidence. The pivotal step in this study is to establish the operating domain prior to aggregation. Pixels outside this domain are neither denoised nor interpolated and are not recorded as weak evidence. This approach constitutes a more focused claim than that of global image enhancement, and this focus is precisely the objective.

The three measurement streams serve distinct, non-interchangeable functions. Autofluorescence provides the redox-sensitive intensity signal and serves as the denominator in ratio calculations. QPI offers a co-registered optical path-length map associated with the specimen structure, but it cannot be reduced to NAD(P)H or FAD intensity. FLIM provides a lifetime derived benchmark that is sensitive to the fluorophore’s binding state. Prior correlative SLAM-GLIM research^38^ demonstrated the value of integrating nonlinear autofluorescence with quantitative phase imaging for morpho-chemical tomography. FPhaS diverges from this approach by using phase: a fixed phase-autofluorescence registration determines the validity of intensity-based redox inference, rather than treating phase as an additional, complementary morphological channel. This fixed registration relies on the shared-objective geometry: because phase and autofluorescence traverse the same objective and stage, their spatial correspondence remains sufficiently stable to establish a permanent reference, ensuring that structural support is defined at the same location as the metabolic signal rather than inferred across modalities that utilize separate optics.

Our validation deliberately maintains a narrow scope of claims, justified by the controls employed. Within cellular regions that are mask-locked, confidence-based selection demonstrated enhanced agreement with FLIM-derived redox estimates. Conversely, for the full field intensity endpoint, the FPhaS confidence map did not surpass the performance of denominator-matched random controls, and the 95% confidence interval included zero. This contrast is interpreted as a specificity control rather than a shortcoming: a general denoising benefit should have manifested at both endpoints, whereas FPhaS improvements are confined to its explicitly defined support domain. This distinction indicates a support domain effect rather than image enhancement. Once cell presence, ratio validity, and confidence weighting are decoupled, pixels deemed unreliable lose the status they previously held alongside quantitatively supported pixels.

That same specificity characterizes the failure modes. A fluorescence-bright pixel with weak phase contrast, for instance, in a thin protrusion or a region of low refractive index contrast, may be excluded even though its fluorescence signal is genuine and measurable. Conversely, a structurally convincing pixel with a weak autofluorescence denominator derives no benefit from the phase map. FPhaS is designed to prioritize specificity over apparent coverage. Fluorescence-only analyses remain entirely feasible outside the support domain; however, such results should be reported without the corresponding support claim.

Support-aware logic remains significant even when pixels are aggregated into cells. MCI, MOI, and MRI preserve information about burden, organization, and reliability that is lost when using whole-cell mean redox ratios. In the dataset comprising 10 fields across 5 A549 cell samples, the descriptor space effectively sorted cells into consistent within-field patterns, although transferability between fields remained limited. A conservative interpretation is appropriate: these descriptors offer support for preserving summaries of cell-level variation rather than a universal phenotype taxonomy. To establish stable biological states associated with the descriptor clusters, perturbation studies, additional cell lines, and larger, independent cultures are necessary.

The reconstruction analysis extends the same principle to acquisition design. Combined WLS improved support-layer metrics with reduced fluorescence sampling, but the reduced acquisition still provides metabolic information, and QPI serves only as an edge-aware structural guide rather than a metabolic surrogate. The defensible claim is correspondingly bounded: within the declared support layer, fluorescence-derived readouts can be better preserved under an eightfold reduction in acquired fluorescence content per channel. Translating that into saved wall clock time, reduced photobleaching, lower phototoxicity, and longer longitudinal viability is a task for prospective studies.

## Conclusion

FPhaS reframes label-free redox imaging around a central question: not what optical redox ratio can be displayed at every pixel, but where that ratio is quantitatively supported. By linking quantitative phase imaging and SLAM autofluorescence through a fixed, experimentally calibrated co-registration, the framework uses phase-derived structure as fluorescence-independent support for interpreting metabolic signal at the same location. This design separates three decisions that are often collapsed into a single segmentation mask: where a cell is present, where a ratio can be formed, and where the estimate has sufficient joint support for quantitative pooling. Reporting these decisions as nested support domains makes the measurement’s operating domain explicit rather than presenting a complete-looking redox map with uneven reliability. Within mask-locked cellular regions, confidence-based selection improved agreement with the FLIM-derived redox endpoint, whereas full-field summaries did not show the same gain, supporting the interpretation that FPhaS improves inference by defining admissible support rather than by globally denoising the image.

Beyond the redox ratio itself, FPhaS provides a principled framework for organizing multimodal evidence. Autofluorescence supplies the redox-sensitive metabolic readout, co-registered phase structure provides fluorescence-independent cellular support, and lifetime imaging serves as an independent validation endpoint. The same support logic extends to cell-level descriptors of metabolic content, metabolic–structural organization and measurement reliability, as well as to reconstruction under reduced fluorescence acquisition burden. These results should be interpreted within their current experimental scope: the present study uses a single cell line, and the proposed descriptors are support-preserving quantitative summaries rather than a fixed biological taxonomy. Broader generality across cell types, perturbations and tissue contexts remains to be established. The main contribution of FPhaS is therefore not a more complete redox image, but an explicit and auditable account of where label-free redox images support quantitative inference.

## Materials and methods

### Multimodal imaging system and cell preparation

FPhaS integrates quantitative phase imaging (QPI), simultaneous label-free autofluorescence multiharmonic (SLAM) microscopy, and fluorescence lifetime imaging microscopy (FLIM)^9-16^ on a unified objective and sample stage (ASI FTP-2000; Applied Scientific Instrumentation, Eugene, OR, USA). Each modality was obtained using the same air objective (CFI75 LWD 20XW; Nikon, Tokyo, Japan) at 20× magnification and 0.8 numerical aperture, thereby minimizing field-level discrepancies among phase, autofluorescence, and lifetime measurements.

QPI used an LED array (64 × 64 RGB LED Matrix; Adafruit Industries, New York, NY, USA) with differential phase contrast illumination at 600 ± 30 nm, which was reconstructed into an optical path length map. For SLAM^9-16^, a femtosecond fiber laser source provided broadband excitation for simultaneous label-free nonlinear imaging, and the emitted nonlinear signals were spectrally split into four detection channels: NAD(P)H (450 ± 30 nm bandpass), FAD (580 ± 30 nm), SHG (515 ± 20 nm), and THG (385 ± 20 nm).

SLAM images were acquired at a resolution of 512 × 512 pixels, with a field of view of 100 μm (corresponding to 0.195 μm per pixel), utilizing a dwell time of 1.8 μs per pixel and averaging over 10 frames. Quantitative Phase Imaging (QPI) images encompassed a 490 μm field at a resolution of 0.109 μm per pixel. Fluorescence Lifetime Imaging Microscopy (FLIM) was performed using the same nonlinear arm, employing time-correlated single-photon counting (PicoHarp 330; PicoQuant, Berlin, Germany) with 8 ps temporal binning, and subsequently analyzed through phasor analysis and least-squares decay fitting. For analysis, one imaging unit was defined as a complete co-registered QPI, SLAM and FLIM image set acquired from a single field of view under one photon-budget condition. The primary A549 cell line validation dataset included 10 fields of view across 5 samples; analyses under high and low-photon conditions were conducted as paired photon-budget conditions when derived from the same cell or field.

FLIM was incorporated into the workflow solely as an orthogonal, lifetime-derived validation readout, never as a component of the FPhaS confidence maps. The fluorescence lifetimes of NAD(P)H distinguish the free species (∼0.4 ns) from the protein-bound species (∼1-3 ns) and are responsive to enzyme-binding states and the local redox environment rather than to the absolute fluorophore concentration. The redox estimates obtained through FLIM thus provide an independent benchmark against which the confidence weighted intensity estimates can be evaluated within matched cellular support.

### Cell culture and live cell imaging

A human alveolar basal epithelial cell line (A549) served as the validation line for these studies. This cell line is derived from lung adenocarcinoma and has been widely used in pulmonary, cancer, and toxicology research^39,40^. Cells were maintained in RPMI-1640 medium containing glucose, L-glutamine, and HEPES, supplemented with 10% fetal bovine serum and 1% penicillin-streptomycin-amphotericin, and imaged at passage 3 in maintenance medium. For imaging, cells were seeded to reach a subconfluent density at acquisition of about 1 × 10□ cells cm□^2^, sparse enough that QPI could resolve individual cell boundaries.

Live-cell measurements were carried out on the microscope stage of a Tokai Hit STXG-WSKMX-SET stage-top incubation system (Tokai Hit, Shizuoka, Japan). The incubator held 37 °C, a humidified atmosphere, and 5% CO□ throughout imaging, and QPI, SLAM, and FLIM were all acquired under this one multimodal geometry.

### One-time multimodal co-registration calibration

Co-registration was calibrated utilizing paired QPI and SLAM images of a USAF 1951 target, with 61 landmark correspondences^22^ distributed across the field of view. For the *ith* landmark, let *p*_*i*_ represent the landmark in the SLAM coordinate system and *q*_*i*_ the corresponding landmark in the QPI coordinate system. The candidate transformations *T* encompassed affine, projective, second-order polynomial, and third-order polynomial mappings. Since the USAF 1951 target is nonfluorescent, we employed the third-harmonic generation signals produced at its chrome glass edges as the SLAM side landmarks, matching them against the edges derived from QPI of the same features, thereby ensuring that the calibration mapping accurately reflected the highly nonlinear signal path employed during biological imaging.

For the selected second-order polynomial model, the mapping from SLAM to QPI coordinates was:

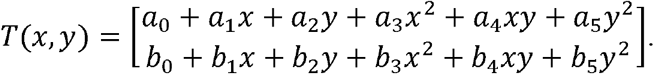

The transform coefficients were estimated from the calibration landmarks by least squares.

For each landmark, the target registration error *e*_*i*_ was computed as:

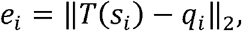

and the root-mean-square residual was

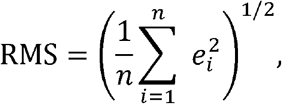

with n = 61. For model selection, we drew on several lines of evidence at once: in-sample residuals, leave-one-out error, random split cross validation, shifted block cross validation, Jacobian regularity, and inverse consistency^22^.

For the inverse consistency analysis, we evaluated the forward mapping *T* and the inverse mapping *T*^−1^ through their round-trip error,

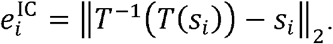

Local geometric regularity was read off the Jacobian matrix (ref. 19),

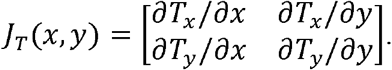

Pixels with a positive Jacobian determinant^23^ were regarded as locally orientation preserving. The second-order polynomial transformation was chosen for its balance between registration precision and geometric stability. Once determined, the transformation was fixed and uniformly applied to all biological images. No biological field was utilized to refine the mapping. The convex hull of the calibration landmarks delineated the validated interpolation domain for pixel level analyses.

### Background correction and image preprocessing

Before any downstream analysis, the QPI, NAD(P)H, and FAD images were background correction. For each channel image *I*(*x*), noncellular pixels provided an estimate of a smooth background field *B*_*I*_ (*x*), and the corrected image was:

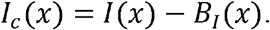

Gaussian smoothing widths were set to 60 pixels for QPI and to 40 pixels for NAD(P)H and FAD. Direct estimation required at least 80 background pixels; in cases with fewer pixels, a resilient fallback at the field level was employed. Background noise was quantified using the residual non-cellular pixels through median absolute deviation.

NAD(P)H, FAD, and QPI backgrounds were estimated independently. After correction, negative autofluorescence values were not used as a valid photon-supported signal for support domain construction. For ratio and confidence calculations, the corrected NAD(P)H and FAD intensities were clipped to non-negative values

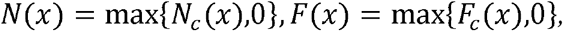

where *N*(*x*)and *F*(*x*)are the processed NAD(P)H and FAD autofluorescence intensities.

Cell segmentation was performed on the QPI images using a Cellpose ^41^ cytoplasmic model. The decision to segment on QPI was intentional: it eliminates the circular dependency that would otherwise couple the cell mask to autofluorescence intensity. The QPI image was normalized comprehensively prior to analysis. The Cellpose^41^ diameter parameter was set to 120 pixels, the cell-probability threshold to 0, and the flow threshold to 0.4. Subsequently, objects smaller than 500 pixels were excluded.

Let *M*_QPI_(*x*) ∈ {0,1} be the binary QPI-derived cell mask; the corresponding QPI cell domain was then

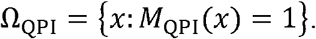

To reduce boundary mixing, we stripped a two-pixel rim from this domain before any ratio or confidence calculation:

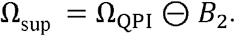

Here *B*_2_is a disk-shaped structuring element of radius two pixels, and denotes binary erosion. Equivalently,

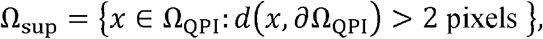

where *d*(*x*, ∂ Ω_QPI_) is the Euclidean distance from the pixel *x* to the QPI mask boundary.

### Optical redox ratio and ratio-of-sums pooling

The total autofluorescence support image was defined as

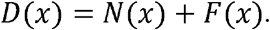

The intensity optical redox ratio^5–7^ was

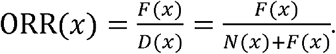

Cell level and domain level redox estimates were obtained using ratio-of-sums pooling. For a pixel domain Omega with nonnegative weights *w*(*x*), the pooled redox estimate was

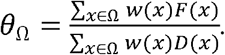

This pooling rule preserves numerator and denominator support before division. It was used instead of averaging local ratio values after division.

Pixelwise ORR uncertainty was approximated by first-order error propagation^27,28^. The full propagated uncertainty model appears in Supplementary Methods. In brief, the target ORR uncertainty was fixed at

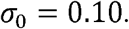

The denominator support floor was

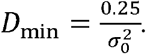

For the final configuration,

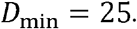

We applied this threshold as a measurement support rule, not as a display threshold.

### Support domain construction

FPhaS defines three nested support domains. The structural domain keeps the QPI supported cell pixels whose phase derived structural reliability is high enough:

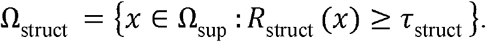

The physical ratio domain retained the pixels whose denominator support is sufficient and whose propagated ORR uncertainty is below the target threshold:

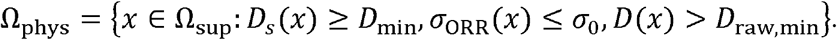

Here *D*_*s*_(*x*) is the smoothed denominator support image, and *D*_raw,min_ =1. The hard operating domain was

Pixels lying outside Ω_hard_ were excluded from the primary quantitative summaries. We did not impute them, recover them by interpolation, or allow display smoothing to fold them back into the analysis.

### Confidence-field construction

The FPhaS confidence field combined fluorescence support with QPI derived structural support. A physical reliability score R_phys_(*x*) summarized the denominator support and propagated ORR uncertainty. A structural reliability score *R*_struct_(*x*) summarized phase derived mass support, interior support, local gradient structure, and local texture. The detailed definitions of both component scores are provided in the Supplementary Methods. In the final configuration, the structural threshold retained the upper 70% of in cell pixels as QPI-supported, and the mask locked *FLIM*_ORR_ agreement remained statistically stable as the cut was swept across the 10th-50th-percentile range (Supplementary Fig. S3c).

The raw FPhaS confidence weight was

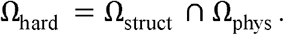

with

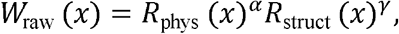

Weights were applied exclusively within the Ω_hard_ operating domain. Non-zero weights

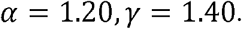

were clipped to the 1st-99th percentiles and subsequently normalized by their median in order to enhance numerical conditioning. The final confidence weight, denoted *W*_(x)_, was set to zero everywhere outside the hard operating domain.

The FPhaS intensity ORR^27,28^ estimate for the cell *k* was

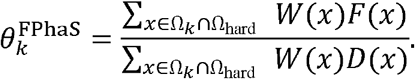

The unweighted estimator used the same domain as *W*(*x*). For a weighted measurement over domain Omega, the effective sample size^31^ was

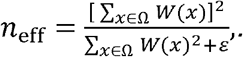

The effective-sample-size fraction was

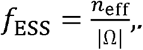

Per-cell quality-control summaries recorded cell area, admissible fraction, denominator support, mean confidence, and effective-sample-size fraction.

### Channel complementarity analysis

We compared the co-registered QPI, NAD(P)H, FAD, and derived fluorescence channels in three ways: marginal Pearson correlation, partial correlation after linear de-confounding, and principal component analysis. For two channels X and Y, the marginal Pearson correlation was

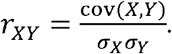

The control-channel set Z contains the variables held constant during partial-correlation estimation^25,26^. We set Z = {NAD(P)H, FAD} when measuring the residual QPI versus fluorescence association, and Z = {QPI mass proxy, QPI gradient} when measuring the residual NAD(P)H versus FAD association. Partial correlation^24^ was computed after removing the linear effects of a control-channel set *Z*. Letting *X*^⊥^ and *Y*^⊥^ denote the residuals after regression on *Z*, the partial correlation was

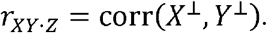

The Pearson to partial attenuation was summarized as

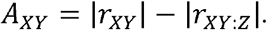

Principal component summaries^25,26^ were subsequently used to assess whether the multimodal stack converged on a single intensity axis.

### FLIM-referenced validation

FLIM derived redox information was used as an orthogonal validation readout. Estimators were compared inside a shared mask locked validation domain,

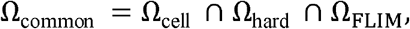

where Ω_FLIM_ denotes the pixels with valid FLIM-derived redox measurements. This shared domain design ensured that every estimator was evaluated on the same cellular support.

Component maps derived from FLIM were acquired through phasor analysis^17–20^: *F*_FLIM_(*x*) represents the *FLIM*_FAD_ map (the bound-fraction-weighted FAD intensity from phasor decomposition), and *D*_FLIM_ is the corresponding denominator-support map, defined as the sum of *F*_FLIM_(*x*) and the bound-fraction-weighted NAD(P)H intensity, ensuring that the

*FLIM*_*ORR*_ retains the same *FAD*/(*FAD*+*NAD*(*P*)*H*) form as the intensity ORR. When these maps were available, the mask-locked FLIM-derived ORR followed from ratio-of-sums pooling^7^:

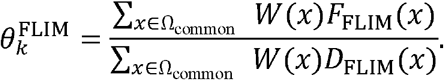

For an intensity derived estimator 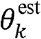, the absolute error against the FLIM derived endpoint was

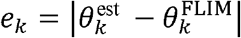

A mask lock audit verified that the FLIM derived redox values were computed within the cell masks supported by QPI. Endpoint analyses were conducted separately for the intensity ORR and the *FLIM*_*ORR*_.

Denominator matched random controls^30^ were employed to evaluate whether the observed confidence based improvements could be solely attributed to the selection of denominators. For each cell and endpoint, random control sets were sampled from the identical candidate domain, ensuring an exact match in both the denominator-support distribution and the coverage of the primary confidence selected pixels. Subsequently, the endpoint was recalculated for each matched random set utilizing the same ratio-of-sums rule. The control statistic was

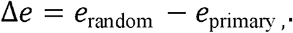

A positive Δ*e* indicates that the primary FPhaS confidence map exhibited a lower error rate compared to the denominator-matched random selector.

### High- and low-photon per pixel conditions

Photon support robustness was evaluated under high and low photon per pixel conditions, with the former corresponding to higher photon counts per pixel and the latter to lower counts. We used these conditions to test whether confidence ranking, *FLIM*_*ORR*_ agreement, and the cell level descriptors remained stable as photon support decreased.

These photon support conditions differ from the spatial binning used in the reconstruction branch. Photon support conditions test robustness to changes in the number of photons per pixel, whereas spatial binning changes the effective sampling grid.

### Cell-level descriptor analysis

Cell level descriptors were computed from confidence weighted FPhaS measurements. The QPI derived dry-mass proxy^15,16^ was

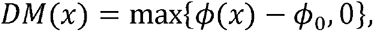

where *ϕ*_0_ is the background phase baseline.

Mass-normalized descriptors used an effective mass image:

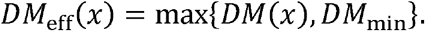

For the cell *k*, the weighted redox endpoint was

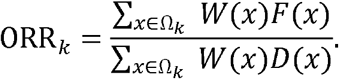

The metabolic content index (MCI) for channel Y was

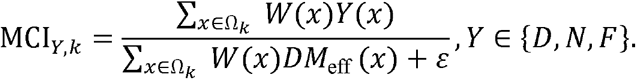

The metabolic reliability index (MRI) was

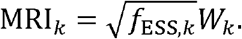

where

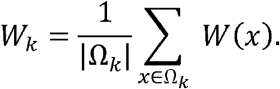

The metabolic organization index (MOI) was established as a collection of structure-metabolism coupling metrics, including local metabolic-structural concordance, structure-conditioned ORR contrast, and the mutual information between QPI dry mass and ORR. Comprehensive definitions of the descriptors, along with permutation-null tests^30^ and analyses of descriptor stability, are available in the Supplementary Methods.

Descriptor stability was evaluated between the high and low photon conditions across different fields of view and cells. The spatial-state maps were considered as image derived organizations of the sampled fields rather than as fixed biological subtypes.

### Reduced-acquisition reconstruction

The reconstruction branch tested whether fluorescence readouts could withstand a reduced acquisition burden. The low photon condition’s image served as the measured input, and the high photon condition’s image served as the reference for benchmark evaluation. We evaluated reconstructions within the same QPI supported cellular masks used for the support layer analysis.

QPI guided edge weights reduced smoothing across phase defined structural boundaries, consistent with the general principles underlying edge preserving weighted least squares^34^ and guided filtering^35^.

Let *y* be the reduced fluorescence observation, *x* is the reconstructed fluorescence image, P is the bloc mean forward operator, and φ denotes the co-registered QPI guide. CombinedWLS reconstructed *x* by minimizing the binning aware weighted least squares objective.

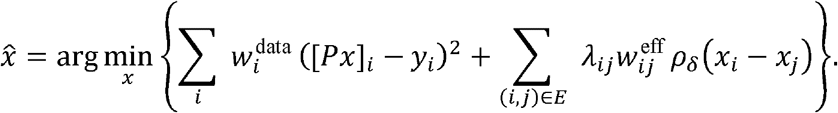

Here *E* is the four-neighbor graph over the support mask, 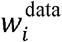 is the photon support data weight, 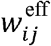 is the QPI guided edge weight, *λ*_*ij*_ is the local regularization strength and *ρ*_*δ*_ is the Huber^42^ penalty. The final configuration used delta = 500.

CombinedWLS was solved using three iteratively reweighted least squares iterations and a conjugate gradient solution^43^ of the resulting linear systems. The conjugate gradient tolerance was 10□□, with a maximum of 800 iterations. When a channel required it, non-negative reconstructions were clipped after each solve.

The main benchmark compared the reduced input, guided filtering, CombinedWLS, and the oracle upper bound within the same support layer evaluation domain. Legacy interpolation and Richardson-Lucy^36,37^ controls, calibration variants, and guide-use controls are reported in the Supplementary analyses.

### Reduced-acquisition comparison

For the acquisition-burden analysis, we conducted a comparison between the high photon condition reference and the eightfold low photon condition acquisition burden within the same support layer evaluation framework. The low photon condition acquisition condition recorded less fluorescence information while maintaining the same cellular support used for reconstruction assessment. The acquisition time reduction factor was

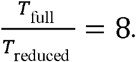

Here *T*_full_ and *T*_reduced_ are the relative fluorescence acquisition content for the high photon condition reference and low photon condition acquisition conditions. We report this as an acquisition burden reduction rather than a direct exposure-time reduction unless prospective timing logs are available. *T*_full_ is the product of pixel dwell time, pixel count, and the number of accumulated frames per channel under the high photon condition acquisition protocol; *T*_reduced_ is the same product under the low photon condition acquisition protocol, with eightfold lower spatial sampling of the fluorescence channels.

Image quality after CombinedWLS reconstruction was scored using the same support layer metrics (detrended NRMSE, Pearson correlation, SSIM^44^, and edge-preservation index). Preservation of the intensity based metabolic readouts was evaluated against the high photon condition acquisition reference within the same support mask.

### Reconstruction metrics

Reconstruction quality was evaluated on support, core, and full region masks. The support mask covered the QPI-supported cellular region. The core mask used an eroded cell region to suppress boundary effects. The full-region mask covered the broader QPI-supported field.

Before evaluation, fluorescence images were detrended by subtracting a Gaussian-fluorescence reference and Ωthe evaluation mask. The root-mean-square error was smoothed background. Let 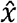 be the reconstructed image, *x*_ref_ the registered high resolution fluorescence reference and Ω the evaluation mask. The root-mean-square error was

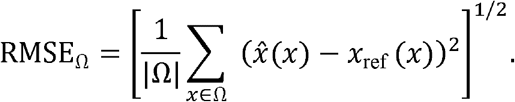

The normalized RMSE was

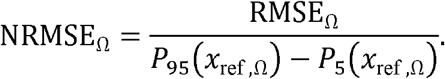

Pearson correlation was computed over the finite masked pixels:

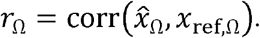

Structural similarity was measured using the SSIM^44^ index. The edge preservation index was the correlation between the Sobel gradient magnitudes of the reconstruction and the reference, computed over the masked pixels:

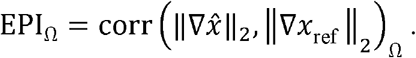

Block artifacts were quantified using a study defined boundary to non-boundary gradient ratio, and frequency resolved agreement was evaluated using Fourier ring correlation. Full metric definitions are provided in the Supplementary Methods.

### Statistical analysis and reproducibility

Registration accuracy (Fig. 1c) is reported as RMS residuals with leave-one-out and random-split cross-validation; these are descriptive error metrics rather than hypothesis tests. Inter-channel associations (Fig. 1e–g) were assessed using Pearson and partial correlations. Confidence intervals for the validation and descriptor endpoints were estimated using a field-of-view cluster bootstrap. Denominator-matched random controls (Fig. 3b) were compared by paired error differences with bootstrap 95% confidence intervals. Descriptor-state associations (Fig. 5g) were assessed by permutation testing. Where significance markers are shown, *p < 0.05, **p < 0.01 and ***p < 0.001. All randomized procedures used fixed seeds recorded in the analysis configuration.

## Code availability

Custom code for registration, support domain construction, confidence field computation, MCI/MOI/MRI descriptor extraction and CombinedWLS reconstruction is available on GitHub and will be archived on Zenodo with a permanent DOI upon publication.

## Data availability

Processed source data underlying the figures and quantitative claims, along with the calibration landmark set, representative QPI/SLAM/FLIM cell stacks, and per-cell descriptor tables, will be deposited on Zenodo with a permanent DOI before publication. Raw imaging data from the studies reported here are available upon reasonable request from the corresponding author.

## Acknowledgements

The authors thank Darold Spillman Jr. for administrative support. This work was supported in part by the U.S. National Institutes of Health (R01EB028615, P41EB031772, T32ES007326) and the Paul G. Allen Frontiers Group.

## Author contributions

H.F. and Z.Y. conceptualized the project. H.F., J.S. and Z.Y. built the co-registered microscope and developed the instrument-control components. H.F. performed the biological experiments. H.F. and J.S. developed the analysis pipelines. A.H., L.Y. and K.K.D.T. provided insights into control-software models. E.A. provided A549 cells and contributed to biological interpretation. H.F. and S.A.B. conceived the project and supervised the work. H.F. wrote the manuscript with input from all authors. S.A.B. supervised and funded the project.

## References

1. Chance,B., Cohen, P., Jobsis, F. & Schoener, B. Intracellular Oxidation-Reduction States in Vivo. Science. 137. 499–508 (1962).

2. Georgakoudi, I. & Quinn, K. P. Optical Imaging Using Endogenous Contrast to Assess Metabolic State. Annu. Rev. Biomed. Eng. 14, 351–367 (2012).

3. Kolenc, O. I. & Quinn, K. P. Evaluating Cell Metabolism Through Autofluorescence Imaging of NAD(P)H and FAD. Antioxidants & Redox Signaling 30, 875–889 (2019).

4. Skala, M. C. et al. In vivo multiphoton microscopy of NADH and FAD redox states, fluorescence lifetimes, and cellular morphology in precancerous epithelia. Proceedings of the National Academy of Sciences 104, 19494–19499 (2007).

5. Cannon, T. M., Shah, A. T., Walsh, A. J. & Skala, M. C. High-throughput measurements of the optical redox ratio using a commercial microplate reader. J Biomed Opt 20, 010503 (2015).

6. Ostrander, J. H. et al. Optical Redox Ratio Differentiates Breast Cancer Cell Lines Based on Estrogen Receptor Status. Cancer Res 70, 4759–4766 (2010).

7. Hu, L., Wang, N., Cardona, E. & Walsh, A. J. Fluorescence intensity and lifetime redox ratios detect metabolic perturbations in T cells. Biomed. Opt. Express, BOE 11, 5674–5688 (2020).

8. Park, J. & Gao, L. Advancements in fluorescence lifetime imaging microscopy Instrumentation: Towards high speed and 3D. Current Opinion in Solid State and Materials Science 30, 101147 (2024).

9. Georgakoudi, I. et al. Consensus guidelines for cellular label-free optical metabolic imaging: ensuring accuracy and reproducibility in metabolic profiling. JBO 30, S23901 (2025).

10. You, S. et al. Intravital imaging by simultaneous label-free autofluorescence-multiharmonic microscopy. Nat Commun 9, 2125 (2018).

11. Boppart, S. A., You, S., Li, L., Chen, J. & Tu, H. Simultaneous label-free autofluorescence-multiharmonic microscopy and beyond. APL photonics 4, 100901 (2019).

12. Boppart, S. A. et al. Label-free optical imaging technologies for rapid translation and use during intraoperative surgical and tumor margin assessment. Journal of biomedical optics 23, 021104 (2017).

13. Sun, Y. et al. Real-time three-dimensional histology-like imaging by label-free nonlinear optical microscopy. Quantitative Imaging in Medicine and Surgery 10, 2177 (2020).

14. You, S. et al. Label-free deep profiling of the tumor microenvironment. Cancer research 81, 2534–2544 (2021).

15. Park, Y., Depeursinge, C. & Popescu, G. Quantitative phase imaging in biomedicine. Nature Photon 12, 578–589 (2018).

16. Popescu, G. et al. Optical imaging of cell mass and growth dynamics. American Journal of Physiology-Cell Physiology 295, C538–C544 (2008).

17. Datta, R., Heaster, T. M., Sharick, J. T., Gillette, A. A. & Skala, M. C. Fluorescence lifetime imaging microscopy: fundamentals and advances in instrumentation, analysis, and applications. JBO 25, 071203 (2020).

18. Lakowicz, J. R., Szmacinski, H., Nowaczyk, K. & Johnson, M. L. Fluorescence lifetime imaging of free and protein-bound NADH. Proceedings of the National Academy of Sciences 89, 1271–1275 (1992).

19. Digman, M. A., Caiolfa, V. R., Zamai, M. & Gratton, E. The Phasor Approach to Fluorescence Lifetime Imaging Analysis. Biophysical Journal 94, L14–L16 (2008).

20. Ranjit, S., Malacrida, L., Jameson, D. M. & Gratton, E. Fit-free analysis of fluorescence lifetime imaging data using the phasor approach. Nat Protoc 13, 1979–2004 (2018).

21. Tian, L. & Waller, L. Quantitative differential phase contrast imaging in an LED array microscope. Opt. Express, OE 23, 11394–11403 (2015).

22. Fitzpatrick, J. M., West, J. B. & Maurer, C. R. Predicting error in rigid-body point-based registration. IEEE Transactions on Medical Imaging 17, 694–702 (1998).

23. Modersitzki, J. Numerical Methods for Image Registration. (OUP Oxford, 2003).

24. Baba, K., Shibata, R. & Sibuya, M. Partial Correlation and Conditional Correlation as Measures of Conditional Independence. Australian & New Zealand Journal of Statistics 46, 657–664 (2004).

25. Pearson, K. LIII. On lines and planes of closest fit to systems of points in space. The London, Edinburgh, and Dublin Philosophical Magazine and Journal of Science 2, 559–572 (1901).

26. Pedregosa, F. et al. Scikit-learn: Machine Learning in Python. J. Mach. Learn. Res. 12, 2825–2830 (2011).

27. Ku, H. H. Notes on the use of propagation of error formulas. J. Res. Natl. Bur. Stand. C 70C, 263–273 (1966).

28. Oehlert, G. W. A Note on the Delta Method. The American Statistician 46, 27–29 (1992).

29. Muir, R. D., Kissick, D. J. & Simpson, G. J. Statistical connection of binomial photon counting and photon averaging in high dynamic range beam-scanning microscopy. Opt. Express, OE 20, 10406–10415 (2012).

30. Phipson, B. & Smyth, G. K. Permutation P-values Should Never Be Zero: Calculating Exact P-values When Permutations Are Randomly Drawn. Statistical Applications in Genetics and Molecular Biology 9,.

31. Kish directly: Kish, L. Survey Sampling. (John Wiley & Sons, New York, 1965).

32. Efron, B. Bootstrap Methods: Another Look at the Jackknife. The Annals of Statistics 7, 1–26 (1979).

33. Hubert, L. & Arabie, P. Comparing partitions. Journal of Classification 2, 193–218 (1985).

34. Farbman, Z., Fattal, R., Lischinski, D. & Szeliski, R. Edge-preserving decompositions for multi-scale tone and detail manipulation. ACM Trans. Graph. 27, 1–10 (2008).

35. He, K., Sun, J. & Tang, X. Guided Image Filtering. IEEE Transactions on Pattern Analysis and Machine Intelligence 35, 1397–1409 (2013).

36. Richardson, W. H. Bayesian-Based Iterative Method of Image Restoration*. J. Opt. Soc. Am., JOSA 62, 55–59 (1972).

37. Lucy, L. B. An iterative technique for the rectification of observed distributions. The Astronomical Journal 79, 745 (1974).

38. Butola, A. et al. Label-free correlative morpho-chemical tomography of 3D kidney mesangial cells. JBO 31, 036501 (2026).

39. Lieber, M., Smith, B., Szakal, A., Nelson-Rees, W. & Todaro, G. A continuous tumor-cell line from a human lung carcinoma with properties of type II alveolar epithelial cells. Int J Cancer 17, 62–70 (1976).

40. Giard, D. J. et al. In vitro cultivation of human tumors: establishment of cell lines derived from a series of solid tumors. J Natl Cancer Inst 51, 1417–1423 (1973).

41. Stringer, C., Wang, T., Michaelos, M. & Pachitariu, M. Cellpose: a generalist algorithm for cellular segmentation. Nat Methods 18, 100–106 (2021).

42. Huber, P. J. Robust Estimation of a Location Parameter. Ann. Math. Statist. 35, 73–101 (1964).

43. Hestenes, M. R. & Stiefel, E. Methods of conjugate gradients for solving linear systems. J. Res. Natl. Bur. Stand. 49, 409–436 (1952).

44. Wang, Z., Bovik, A. C., Sheikh, H. R. & Simoncelli, E. P. Image quality assessment : from error visibility to structural similarity. IEEE Trans. Image Process. 13, 600–612 (2004)..

