## Supplemental methods for "Confidence-supported label-free metabolic imaging with FPhaS fixed phase–autofluorescence microscopy"

### Supplementary Methods

#### Supplementary Method 1. Background correction and QPI-based segmentation

QPI was acquired by LED-array differential phase contrast imaging^1^. QPI, NAD(P)H and FAD images were background corrected before downstream analysis. For each image channel $I(x)$, a smooth background field $B_{I}(x)$ was estimated from non-cellular pixels. The corrected image was defined as:

$I_{c}(x)=I(x)-B_{I}(x)$.

Background noise was estimated from corrected non-cellular residuals $r_{I}(x)$ using the median absolute deviation.

$\sigma_{I}=1.4826\text{ median }\left( \left| r_{I}-median\left( r_{I} \right) \right| \right)$.

For support-domain construction, the corrected autofluorescence intensities were truncated at zero.

$$N(x)=max\left\{ N_{c}(x),0 \right\}, F(x)=max\left\{ F_{c}(x),0 \right\}.$$

#### Supplementary Method 2. Propagated ORR uncertainty and denominator support

We approximated pixelwise ORR uncertainty by delta-method propagation^2,3^. For

$$ORR(x)=\frac{F(x)}{N(x)+F(x)}$$

the Poisson term^4^ used in the implementation was

$$\mathrm{Var}_{P}[ORR(x)]\approx\frac{N(x)F(x)}{D(x)^{3}}.$$

The local derivatives of $F/D$ are

$$\begin{matrix} & \frac{\partial}{\partial F}\left( \frac{F}{D} \right)=\frac{N}{D^{2}}, \\ & \frac{\partial}{\partial N}\left( \frac{F}{D} \right)=-\frac{F}{D^{2}} \end{matrix}$$

Including background-noise terms gave the total variance approximation

$$\sigma_{\mathrm{ORR}}^{2}(x)=\frac{N(x)F(x)}{D(x)^{3}}+\left( \frac{N(x)}{D(x)^{2}} \right)^{2}\sigma_{F}^{2}+\left( \frac{F(x)}{D(x)^{2}} \right)^{2}\sigma_{N}^{2}$$

The propagated ORR uncertainty was

$$\sigma_{\mathrm{ORR}}(x)=\sqrt{\sigma_{\mathrm{ORR}}^{2}(x)}.$$

The target ORR uncertainty was fixed as

$$\sigma_{0}=0.10.$$

The denominator-support floor was

$$D_{\min}=\frac{0.25}{\sigma_{0}^{2}}.$$

For the final configuration,

$$D_{\min}=25.$$

We applied this threshold as a measurement-support rule, not as a display threshold^5^.

#### Supplementary Method 3. QPI structural reliability

The QPI structural reliability score was computed from four phase-derived components: mass support, interior support, low-gradient support and low-texture support^6^. Let $\phi(x)$ denote the background-corrected QPI optical-path-length image. A smoothed phase image was denoted by $\phi_{s}(x)$, and a smoothed mass-support image was denoted by $\phi_{m}(x)7$.

The robust percentile-scaling operator was defined as

$$robust01\left( z \right)=clip\left( \frac{z-P_{1}\left( z \right)}{P_{99}\left( z \right)-P_{1}\left( z \right)},0,1 \right).$$

where the percentile was fixed to the 30th percentile of the in-cell phase-derived structural-reliability distribution. The threshold pool was the physical candidate domain when sufficient physical-support pixels were available; otherwise, the full eroded QPI cell domain was used.

The mass-support term was

$$C_{\text{mass }}(x)=robust01\left( \phi_{m}(x) \right).$$

The interior-support term was

$$C_{\text{int }}(x)=min\left\{ \frac{d\left( x,\partial\Omega_{\mathrm{QPI}} \right)}{d_{0}},1 \right\}, d_{0}=6\text{ pixels. }$$

The low-gradient support term used the gradient magnitude of the smoothed QPI image:

$$C_{\mathrm{grad}}(x)=1-robust01\left( \left\| \nabla\phi_{s}(x) \right\|_{2} \right).$$

The local phase texture was

$$\sigma_{\phi,\text{ local }}\left( x \right)=\left[ G_{\sigma_{t}}\left( \phi_{s}^{2} \right)\left( x \right)-G_{\sigma_{t}}\left( \phi_{s} \right)^{2}\left( x \right) \right]^{\frac{1}{2}}.$$

where $G_{\sigma_{t}}$ denotes Gaussian smoothing at the texture scale. The low-texture support term was

$$C_{\text{tex }}(x)=1-robust01\left( \sigma_{\phi,\text{ local }}(x) \right)$$

The QPI structural reliability score was the weighted geometric mean

$$R_{\text{struct }}(x)=\left[ C_{\text{mass }}(x)^{1.0}C_{\text{int }}(x)^{1.0}C_{\text{grad }}(x)^{0.6}C_{\text{tex }}(x)^{0.4} \right]^{1/3}$$

Values outside $\Omega_{\mathrm{QPI}}$ were set to zero.

In the final configuration, the structural threshold was fixed so that $R_{struct(x)}\geq\tau_{struct}$ retained the upper 70% of in-cell pixels as QPI-supported, with tau_struct set to the 30th percentile of the in-cell phase-derived structural-reliability distribution.

#### Supplementary Method 4. Support-domain construction

FPhaS used nested support domains to separate cell presence, ratio admissibility and confidence-supported quantification.

The structural domain was

$$\Omega_{\text{struct }}=\left\{ x\in\Omega_{\text{sup }}:R_{\text{struct }}(x)\geq\tau_{\text{struct }} \right\}$$

The structural threshold was set as

$$\tau_{\text{struct }}=Q_{0.20}\left( R_{\text{struct }}(x):x\in\Omega_{\text{pool }} \right),$$

where the percentile was fixed to the 30th percentile of the in-cell phase-derived structural-reliability distribution. The threshold pool was the physical candidate domain when sufficient physical-support pixels were available; otherwise, the full eroded QPI cell domain was used.

The physical ratio domain was

$$\Omega_{\mathrm{phys}}=\left\{ x\in\Omega_{\sup}:D_{s}(x)\geq D_{\min}, \sigma_{\mathrm{ORR}}(x)\leq\sigma_{0}, D(x)>D_{raw,min} \right\}.$$

Here $D_{s}(x)$ is the smoothed denominator-support image, and

$$D_{\text{raw },min}=1$$

The hard operating domain was

$$\Omega_{\text{hard }}=\Omega_{\text{struct }}\cap\Omega_{\text{phys }}.$$

Pixels outside $\Omega_{\text{hard }}$ were excluded from primary quantitative summaries. They were not imputed, restored by interpolation or smoothed into the analysis.

#### Supplementary Method 5. Confidence-field construction

The physical reliability field combined denominator support and propagated ORR uncertainty. The sigmoid function was

$$S(z)=\frac{1}{1+exp(-z)}$$

The denominator-support term was

$$R_{\mathrm{den}}(x)=S\left( \frac{D_{s}(x)-D_{\min}}{0.25D_{\min}} \right).$$

The uncertainty-support term was

$$R_{\sigma}(x)=S\left( \frac{\sigma_{0}-\sigma_{\mathrm{ORR}}(x)}{0.03} \right).$$

The physical reliability score was

$$R_{\mathrm{phys}}(x)=\left[ R_{\mathrm{den}}(x)R_{\sigma}(x) \right]^{1/2}.$$

The raw FPhaS confidence weight was

$$W_{\text{raw }}(x)=R_{\text{phys }}(x)^{\alpha}R_{\text{struct }}(x)^{\gamma},$$

with

$$\alpha=1.20, \gamma=1.40.$$

Weights were used only inside $\Omega_{\text{hard }}$. Let $Q_{0.01}$ and $Q_{0.99}$ denote the 1 st and 99 th percentiles of $W_{\text{raw }}$ inside $\Omega_{\text{hard }}$. The clipped weight was

$$W_{\text{clip }}(x)=clip\left( W_{\text{raw }}(x),Q_{0.01},Q_{0.99} \right).$$

The normalized confidence weight was

$$W(x)=\frac{W_{\text{clip }}(x)}{median\left[ W_{\text{clip }}(u):u\in\Omega_{\text{hard }} \right]}.$$

For pixels outside $\Omega_{\text{hard }}$,

$$W(x)=0.$$

#### Supplementary Method 6. High and low-photon conditions

We evaluated photon-support robustness under two regimes. The high-photon condition carried higher photon counts per pixel; the low-photon condition carried fewer. These let us test whether confidence ranking, FLIM-ORR agreement and the cell-level descriptors held steady as photon support fell^8^.

These photon-support conditions are not the spatial binning used in the reconstruction branch. Photon-support conditions probe robustness as the photon counts per pixel change, whereas spatial binning alters the effective sampling grid and entered only the QPI-guided reconstruction.

#### Supplementary Method 7. Cell-level descriptor definitions

The metabolic reliability index was

$$\mathrm{MRI}_{k}=\sqrt{f_{ESS,k}}\bar{W}_{k},$$

where

$$\bar{W}_{k}=\frac{1}{\left| \Omega_{k} \right|}\sum_{x\in\Omega_{k}} W(x).$$

The metabolic organization index was implemented as a family of structure-metabolism coupling measurements. For two maps $A$ and $B$, the weighted mean of $A$ was

$$\mu_{A}=\sum_{x\in\Omega_{k}} \tilde{W}(x)A(x),$$

with normalized weights

$$\tilde{W}(x)=\frac{W(x)}{\sum_{u\in\Omega_{k}} W(u)}.$$

The weighted Pearson correlation between $A$ and $B$ inside cell $k$ was

$$\rho_{w}(A,B)=\frac{\sum_{x\in\Omega_{k}} \tilde{W}(x)\left[ A(x)-\mu_{A} \right]\left[ B(x)-\mu_{B} \right]}{\left[ \sum_{x\in\Omega_{k}} \tilde{W}(x)\left[ A(x)-\mu_{A} \right]^{2} \right]^{1/2}\left[ \sum_{x\in\Omega_{k}} \tilde{W}(x)\left[ B(x)-\mu_{B} \right]^{2} \right]^{1/2}}.$$

Local metabolic-structural concordance was computed in a sliding window $U_{x}$. For maps $A$ and $B$, local correlation was

$$\rho_{\text{local }}(x)=\frac{\mathrm{cov}_{U_{x}}(A,B)}{\left[ \mathrm{var}_{U_{x}}(A)\mathrm{var}_{U_{x}}(B) \right]^{1/2}}$$

The structure-conditioned ORR contrast was

$$\Delta_{\mathrm{DM}}^{\mathrm{ORR}}={\overline{\mathrm{ORR}}}_{W}\left( DM_{\mathrm{eff}}\geq Q_{0.70} \right)-{\overline{\mathrm{ORR}}}_{W}\left( DM_{\mathrm{eff}}\leq Q_{0.30} \right),$$

where $Q_{0.70}$ and $Q_{0.30}$ are cell-specific quantiles of $DM_{\text{eff }}$.

Mutual information between $DM_{\text{eff }}$ and ORR was computed after quantile binning:

$$I(DM;ORR)=\sum_{a,b} p(a,b)\log_{2}\left[ \frac{p(a,b)}{p(a)p(b)} \right]$$

We generated a within-cell permutation null by permuting the ORR values within the same cell support. The co-registration specificity term was

$$COSI=I(DM;ORR)-I\left( DM;\mathrm{ORR}_{\mathrm{perm}} \right)$$

Descriptor stability was assessed between the high and low-photon conditions, across fields of view and under bootstrap resampling. Principal component analysis and clustering were then used to summarize how the descriptors varied across cells. We read the spatial-state maps as image-derived organizations of the sampled fields.

#### Supplementary Method 8. QPI-guided reconstruction

The reconstruction branch used spatially binned fluorescence images as photon-supported low-resolution observations and co-registered high resolution QPI images as structural guides. Spatial binning increased photon support per effective fluorescence pixel but reduced spatial detail. The binning factor was $B=4.$ Spatial binning increased photon support per effective fluorescence pixel but reduced spatial detail, a trade-off common in photon-limited fluorescence and lifetime imaging.

QPI-guided edge weights reduced smoothing across phase-defined structural boundaries, consistent with the established principles of edge-preserving weighted least-squares reconstruction and guided filtering.

Let $y$ denote the photon-supported binned fluorescence observation, $x$ the reconstructed high-resolution fluorescence image and $P$ the block-mean forward operator. The forward operator was implemented as block averaging followed by replication to the high-resolution grid:

$$P(x)=\mathrm{replicate}_{B}\left[ \mathrm{mean}_{B}(x) \right].$$

The co-registered QPI guide was denoted by $\phi$. A four-neighbor graph $E$ was built over the support mask. For neighboring pixels $i$ and $j$, the QPI-guided edge weight was

$$w_{ij}^{\mathrm{phase}}=exp\left[ -\frac{\left( \phi_{i}-\phi_{j} \right)^{2}}{2\sigma_{e}^{2}} \right].$$

The QPI edge scale was

$$\sigma_{e}=2Q_{0.70}\left( \left| \phi_{i}-\phi_{j} \right|:(i,j)\in E \right).$$

The channel-specific coupling factors were fixed from the measured QPI-fluorescence coupling analysis and held constant throughout the final reconstruction branch. The effective edge weight was

$$w_{ij}^{\mathrm{eff}}=c_{m}w_{ij}^{\mathrm{phase}}+\left( 1-c_{m} \right),$$

where $c_{m}$ is the modality-specific coupling factor.

We scaled the data weights by photon-supported signal magnitude, in keeping with the photon-counting noise behavior of fluorescence microscopy. The data weight itself followed from photon support:

$w_{i}^{\mathrm{data}}=\frac{\sqrt{\max\left( y_{i},0 \right)+1}}{\mathrm{mean}_{j\in\Omega_{\sup}}\sqrt{\max\left( y_{j},0 \right)+1}}$.

The normalized local fluorescence support was

$$u_{i}=clip\left( \frac{G_{\sigma}(y)_{i}}{P_{99}\left[ G_{\sigma}(y) \right]+\varepsilon},0,1 \right).$$

where $G_{\sigma}$ denotes Gaussian smoothing.

We adapted the local regularization strength to fluorescence support through a fixed monotonic schedule, which can be expressed as

$$\lambda_{i}=\lambda_{\max}exp\left( -ku_{i} \right)+\lambda_{\min}\left[ 1-exp\left( -ku_{i} \right) \right],$$

with

$$\lambda_{\min}=1, \lambda_{\max}=40, k=5.$$

For graph edge $(i,j)$, the edgewise regularization strength was

$$\lambda_{ij}=\frac{\lambda_{i}+\lambda_{j}}{2}.$$

CombinedWLS was formulated as a binning-aware weighted least-squares reconstruction with edge-preserving QPI-guided^9^ regularization and a Huber penalty^10^. CombinedWLS reconstructed the high-resolution fluorescence image by minimizing the binning-aware weighted least-squares objective

$$\hat{x}=arg\min_{x} \left\{ \sum_{i} w_{i}^{\text{data }}\left( [Px]_{i}-y_{i} \right)^{2}+\sum_{(i,j)\in E} \lambda_{ij}w_{ij}^{\text{eff }}\rho_{\delta}\left( x_{i}-x_{j} \right) \right\}.$$

The Huber loss^10^ was

$$\rho_{\delta}(r)=\left\{ \begin{matrix} \frac{1}{2}r^{2}, & |r|\leq\delta, \\ [4pt]\delta\left( |r|-\frac{1}{2}\delta\right), & |r|>\delta. \end{matrix} \right.$$

The final configuration used $\delta=500$.

CombinedWLS was solved with three iteratively reweighted least-squares iterations and conjugate-gradient solution of the resulting linear systems^11^. The maximum number of conjugate-gradient iterations was 800. Non-negative reconstructions were clipped after each solve when required by the channel^11^.

To bound the mechanism claim of the reconstruction branch, we ran guide-use specificity controls that compared three guide conditions using the same reduced-acquisition input. The no-QPI condition disabled the QPI-guided edge term; the shifted-QPI condition translated the guide by 5 pixels from its calibrated position; and the registered-QPI condition used the frozen FPhaS calibration. These controls confine the QPI guide to serving as an edge-aware regularizer within the declared support layer, rather than as a source of metabolic contrast^12^ (Supplementary Figs. S8 and S9).

#### Supplementary Method 9. Reconstruction metrics

Block artifacts were quantified by a study-defined boundary-to-nonboundary gradient ratio, defined as:

$$\text{ }\text{Blockiness}\text{ }=\frac{mean(|\nabla\hat{x}|\mid\text{ block boundary })}{mean(|\nabla\hat{x}|\mid\text{ nonboundary })+\varepsilon}.$$

Frequency-resolved agreement was evaluated by Fourier ring correlation. For Fourier transforms of two masked images, the Fourier ring correlation at each radial frequency bin was

$$FRC(r)=\frac{\mathrm{Re}\left[ \sum_{k\in r} A(k)^{*}B(k) \right]}{\left[ \sum_{k\in r} |A(k)|^{2}\sum_{k\in r} |B(k)|^{2} \right]^{1/2}}.$$

Numerical values were drawn from the exported notebook outputs and source-data tables^13^. Matched method evaluations used paired comparisons; resampling-based stability estimates used bootstrap intervals; descriptor-state association analyses and within-cell organization nulls used permutation tests^14^. Every randomized procedure used fixed seeds recorded in the analysis configuration.

#### Supplementary Method 10. FLIM-derived ORR components and rationale

FLIM delivers both intensity and lifetime information at every pixel by measuring fluorescence decay kinetics^15^. Phasor analysis^16^ of the FLIM time traces recovers lifetime-resolved component maps, including free and bound NAD(P)H and FAD contributions^17^. In the mask-locked validation, $F_{FLIM(x)}$ denotes the FLIM-derived FAD-associated map, and $D_{FLIM(x)}$ denotes the matched denominator-support map formed from FLIM-derived FAD signal and the corresponding NAD(P)H component. These lifetime-resolved readouts report biochemical contrast related to fluorophore binding^18,19^, enzyme-associated states and the local redox microenvironment; they should not be reduced to a single free/bound interpretation or treated as absolute fluorophore concentration. In FPhaS, these FLIM-derived readouts are deliberately withheld from confidence-map construction and used only as an orthogonal validation endpoint for confidence-weighted intensity ORR estimates within the same mask-locked cellular support domain.

### Supplementary Results and Figures

#### Supplementary Result 1. Registration stress tests support the fixed calibration

Supplementary Fig. S1 expands the registration evidence summarized in main Fig. 1. It shows non-biological USAF landmark selection, local before/after alignment inspection, transform-family comparison, spatial-block cross-validation and rank aggregation^20^. The second-order polynomial transform is retained because it combines subpixel registration error with the most stable behavior under structured holdout, without refitting to biological fields^21^.

**Supplementary Fig. S1 | Registration model selection and fixed-calibration robustness.**

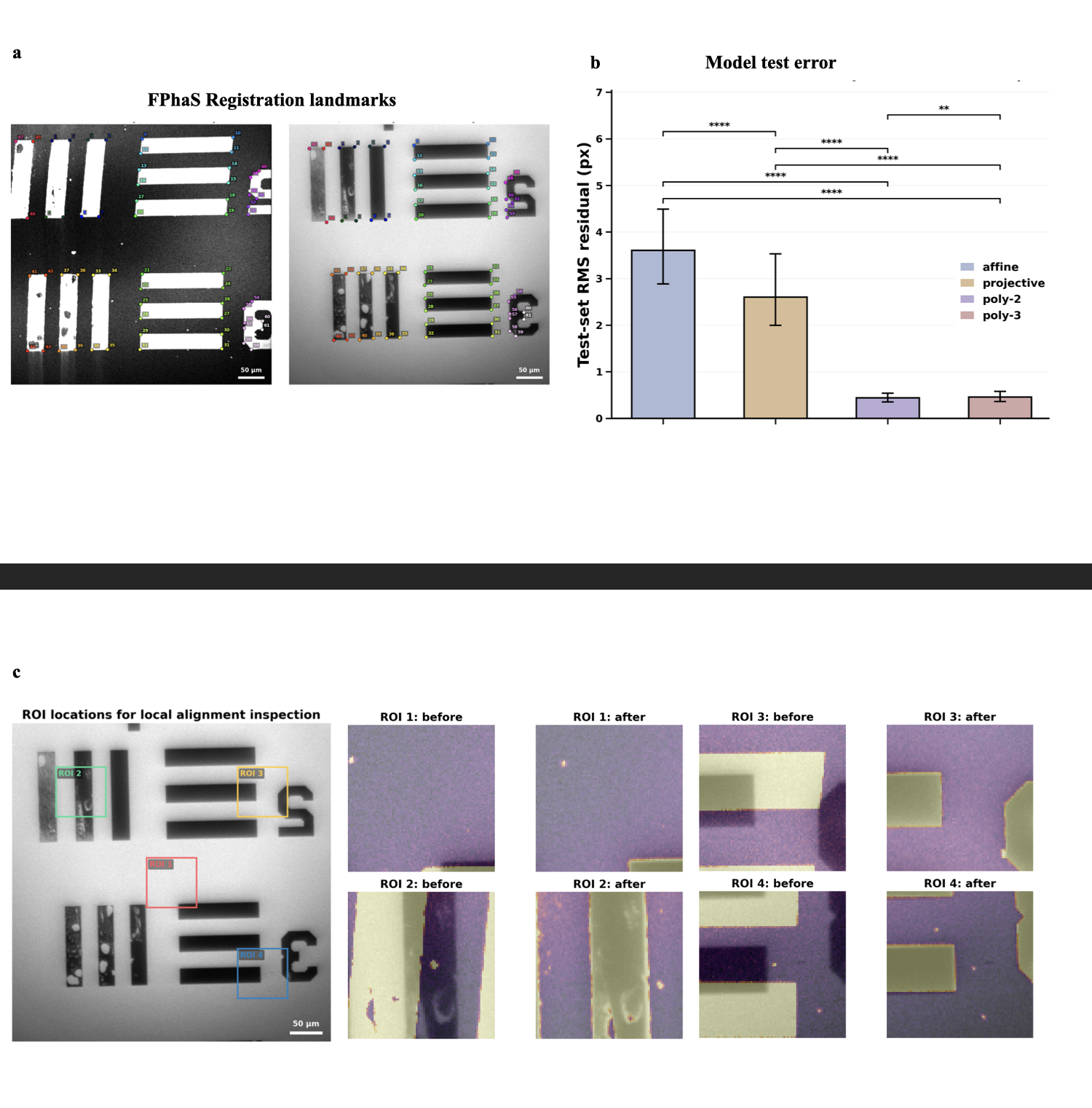

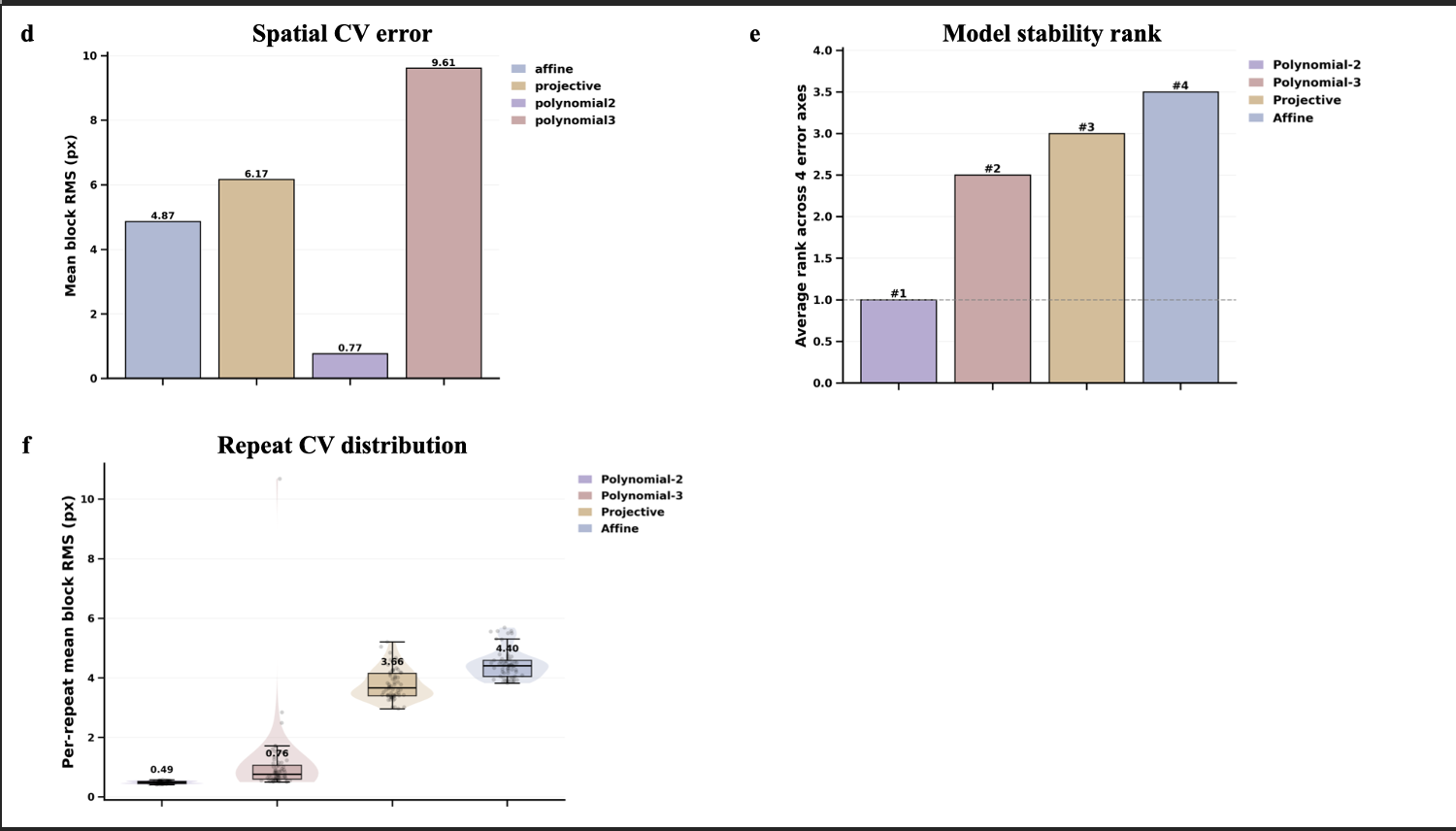

a, USAF 1951 target landmarks used for non-biological QPI-SLAM calibration and b, transform-family comparison. c, Local alignment inspection in four representative target regions before and after applying the retained transform. d, Spatial-block cross-validation, e, stability-rank aggregation, and f, shifted-repeat distribution across candidate transform families. The second-order polynomial calibration was selected from non-biological target data and frozen for all biological fields.

#### Supplementary Result 2. QPI is coupled to but non-redundant with autofluorescence

Supplementary Fig. S2 expands the channel-complementarity analysis. The group-level correlation maps, QPI-coupling summary, partial-correlation matrix, normalized mutual-information matrix and Pearson-minus-mutual-information comparison^22,23^ show that QPI remains coupled to the specimen but is not reducible to NAD(P)H, FAD, harmonic or FLIM-derived channels. These controls justify treating QPI as an external structural support channel rather than a metabolic surrogate^24–26^.

**Supplementary Fig. S2 | Extended QPI-autofluorescence complementarity.**

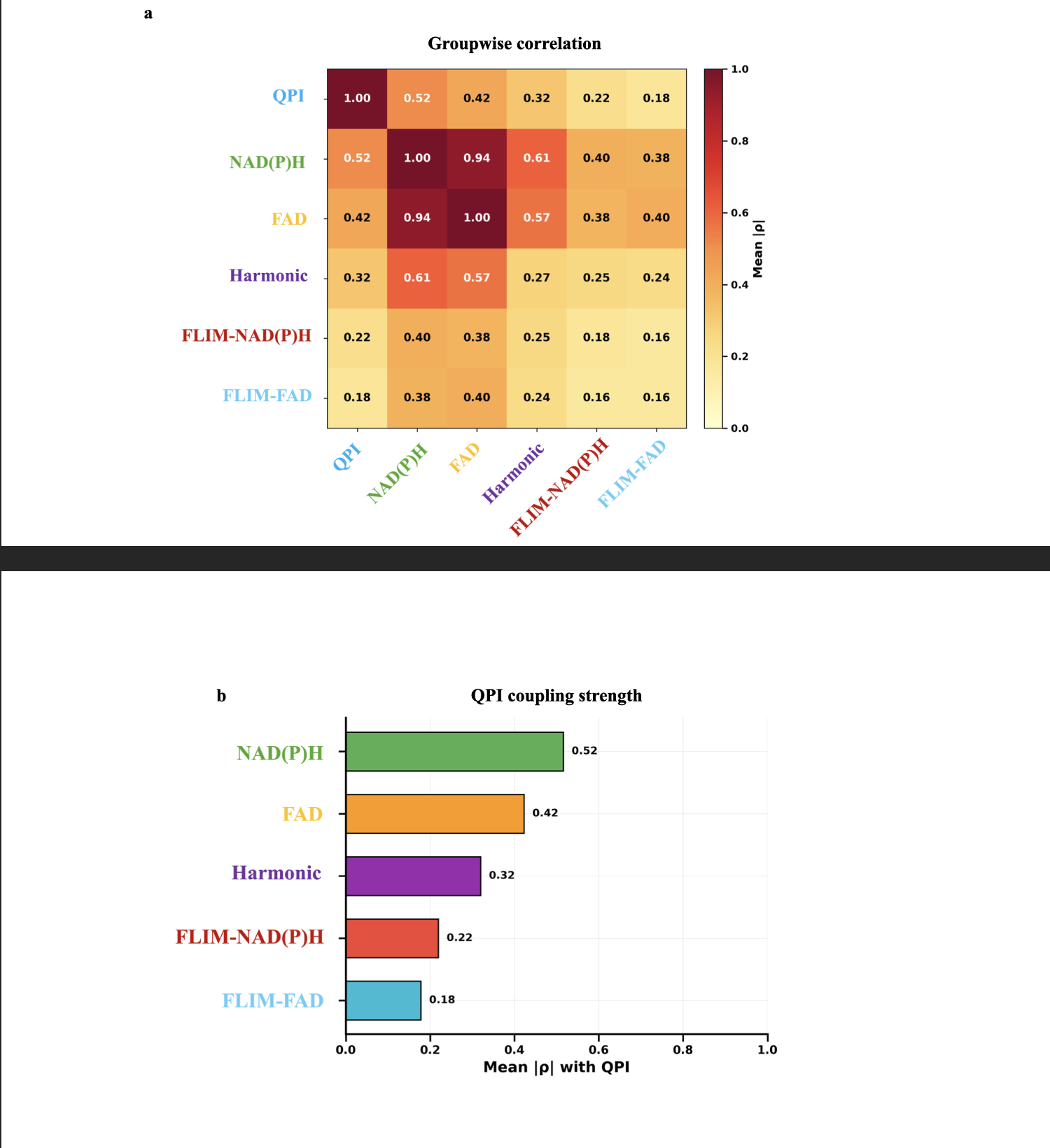

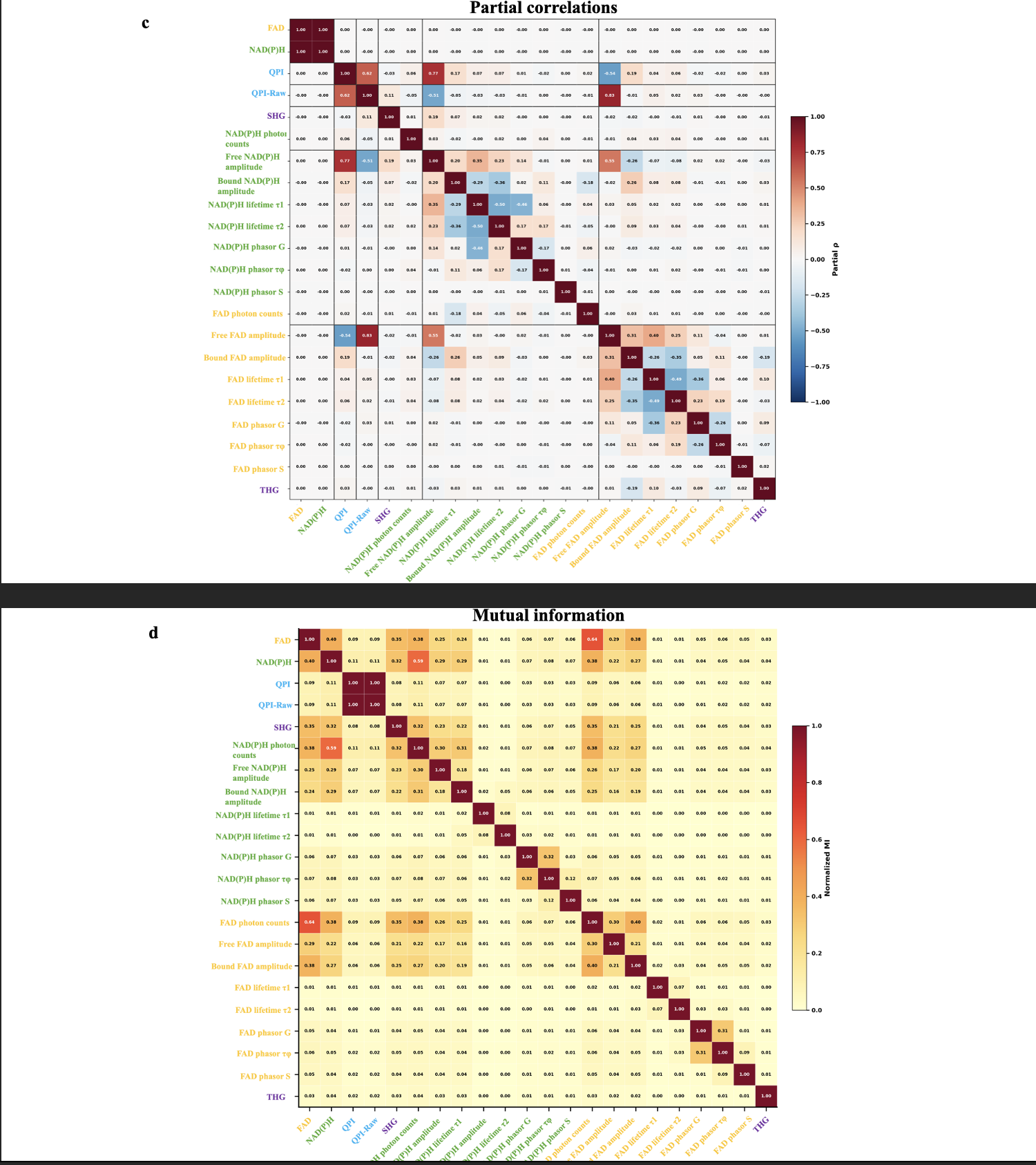

a, Group-level correlation maps among QPI, NAD(P)H, FAD, harmonic-generation and FLIM-derived channel groups. b, Mean QPI coupling strength with each modality group. c, Partial-correlation matrix after de-confounding. d, Normalized mutual-information matrix. Together, these analyses support the use of QPI as a coupled but non-redundant structural support channel rather than a surrogate for autofluorescence or FLIM-derived measurements.

#### Supplementary Result 3. Confidence-selector robustness under photon-support changes

Supplementary Fig. S3 summarizes selector-level robustness under low- and high-photon-per-pixel conditions and compares photon-only, QPI-only, SLAM-only and fused confidence fields. These controls show that support ranking is not a display threshold; it is a measurement-admissibility rule that remains bounded by the FLIM-derived endpoint and the declared cellular support domain.

**Supplementary Fig. S3 | Confidence-selector robustness under photon-support changes.**

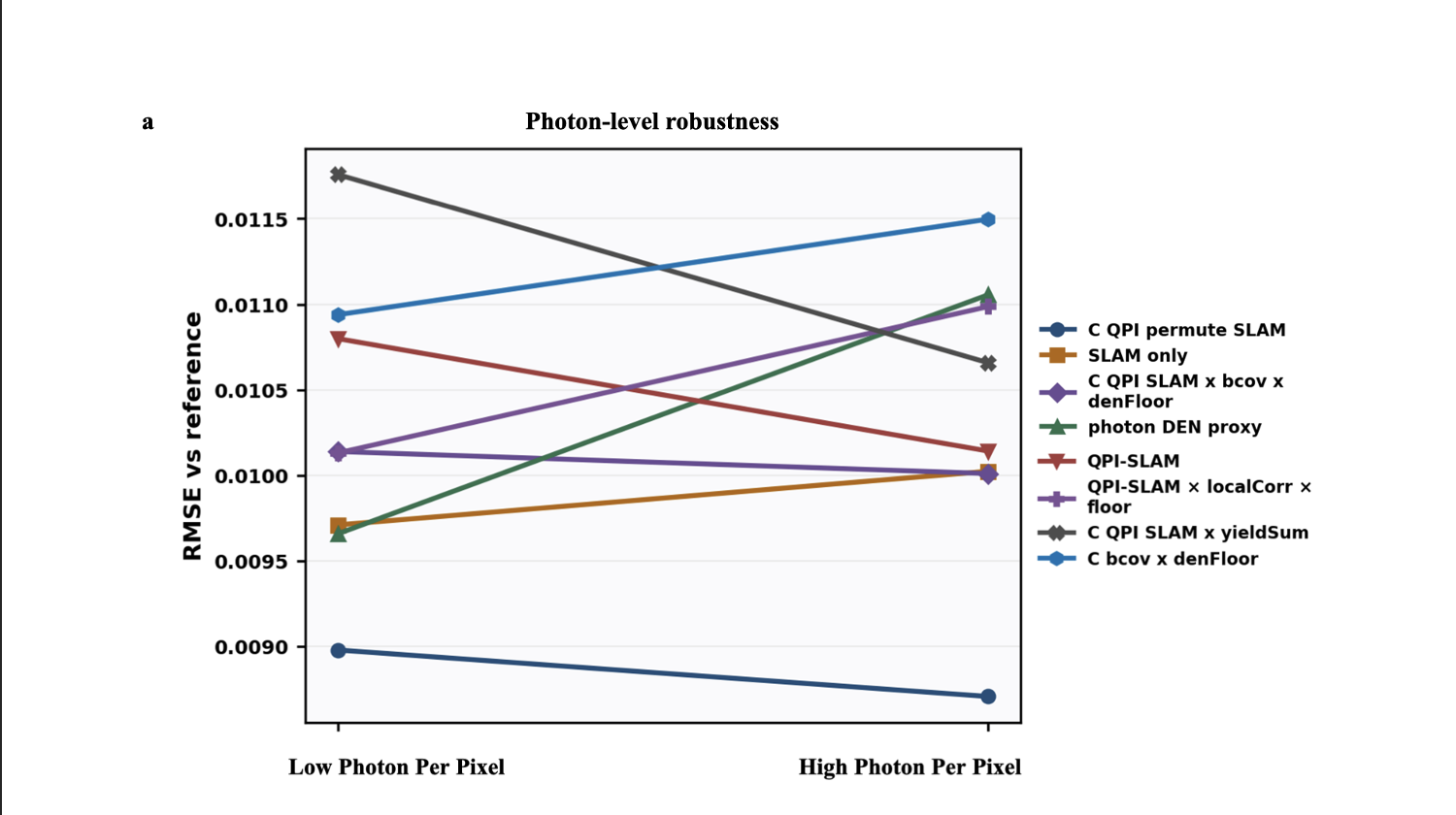

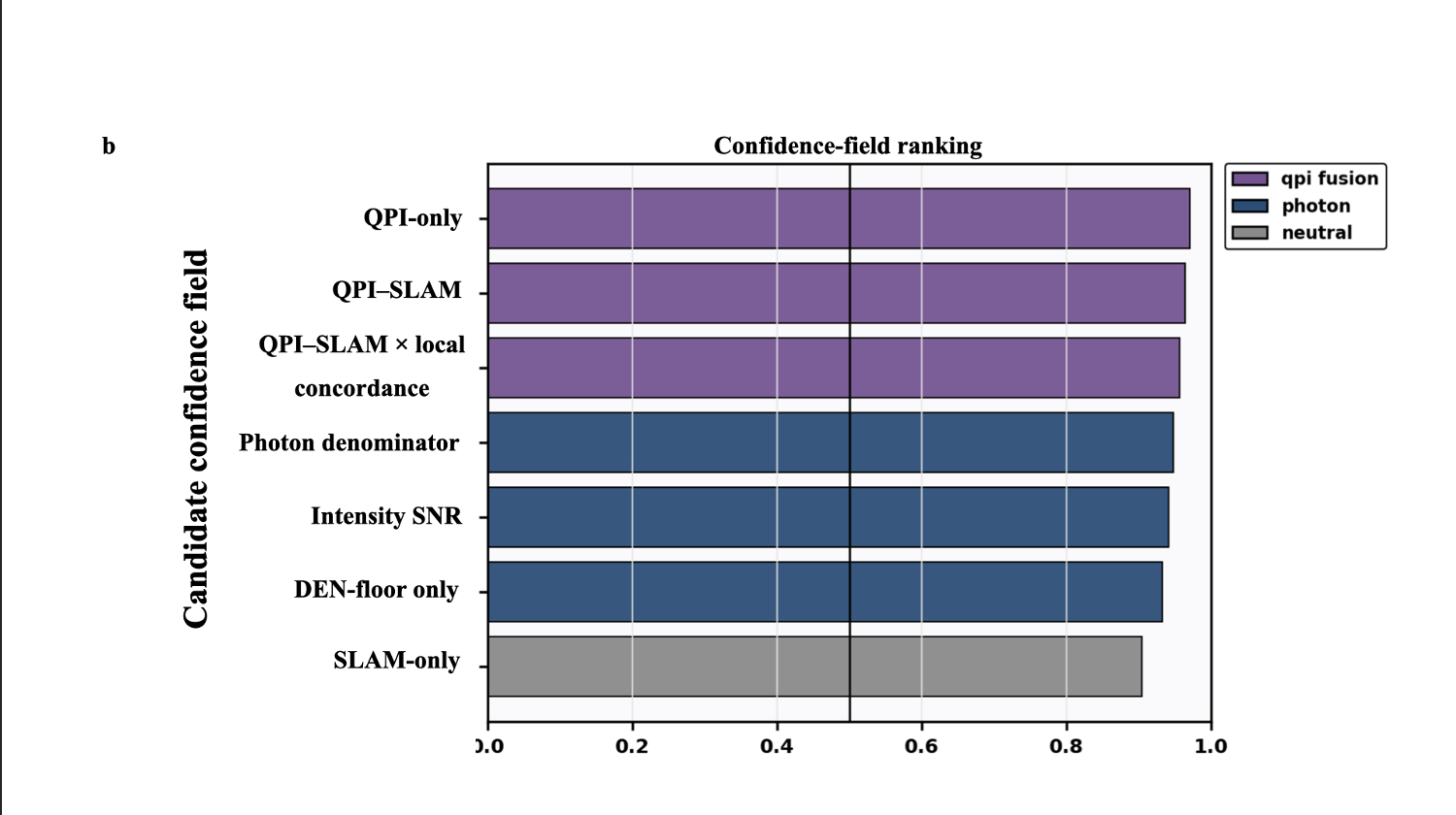

a, shows the selector error relative to the FLIM-derived reference under low and high-photon-per-pixel conditions, and b, shows the confidence-field comparison across QPI-only, QPI-SLAM, fused local-correlation, photon-denominator, intensity-SNR, denominator-floor, and SLAM-only selectors. These controls show that the final support claim is bounded by the declared endpoint and is not equivalent to generic brightness ranking.

#### Supplementary Result 4. Confidence-component maps define the support evidence

Supplementary Fig. S4 shows representative maps for yield consistency, local concordance, biomass covariance confidence, gradient-product concordance and fused confidence products. These component maps clarify how photon support and QPI-derived structural support contribute to the final FPhaS confidence layer while remaining distinct evidentiary sources.

**Supplementary Fig. S4 | Confidence-component maps for support-field construction.**

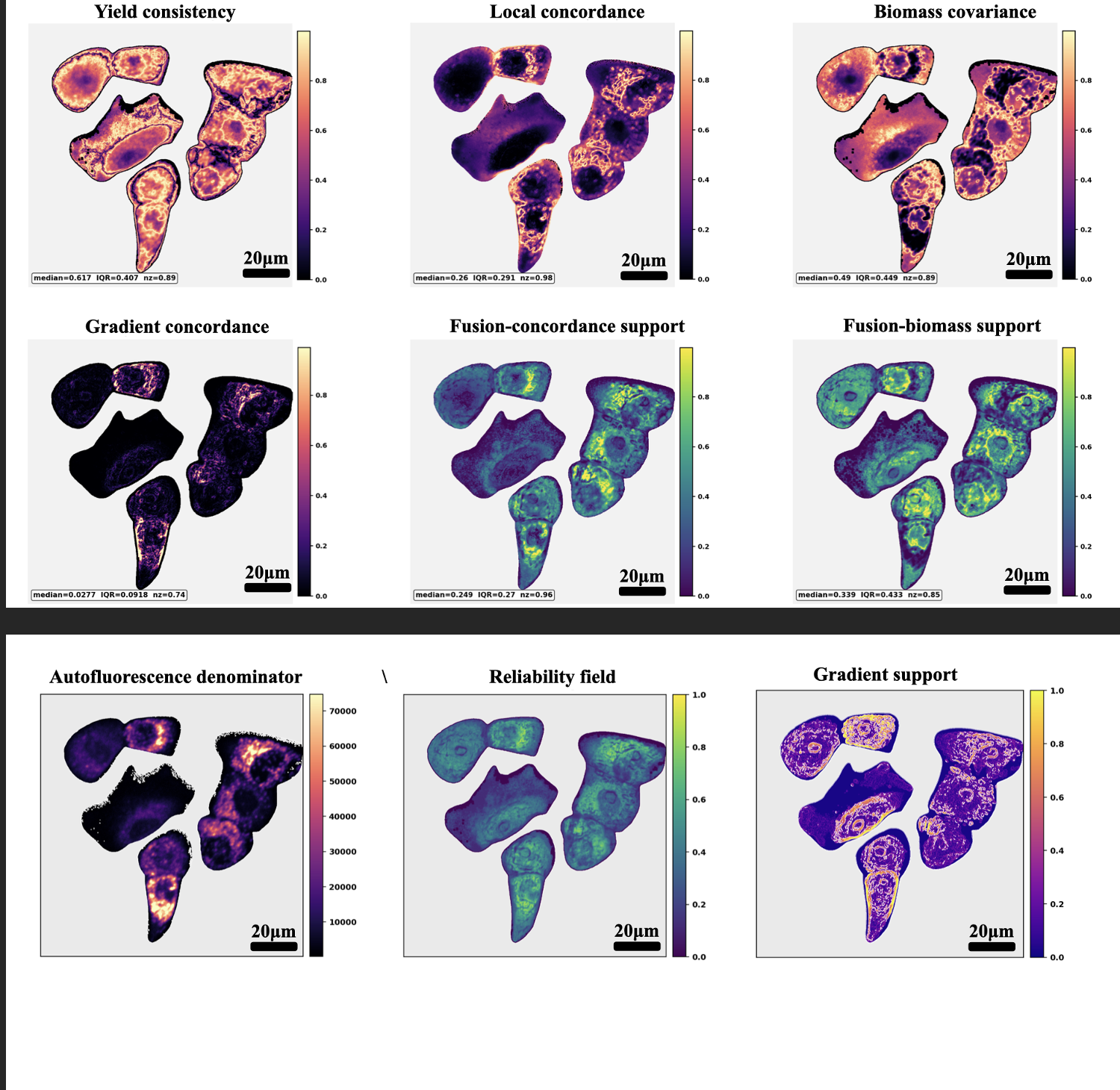

Representative component maps show yield consistency, local concordance, biomass covariance confidence, gradient-product concordance, fusion by local concordance and fusion by biomass covariance, together with denominator, reliability and gradient-support maps. These maps show how photon support and phase-derived structural support contribute spatially distinct information to the FPhaS confidence layer.

#### Supplementary Result 5. FLIM-derived ORR validation is mask-locked and endpoint-specific

Supplementary Fig. S5 provides a representative mechanism case for the held-out FLIM-derived ORR endpoint. It displays QPI dry-mass proxy, SLAM denominator support, intensity ORR, FLIM-derived ORR, absolute intensity-versus-FLIM error, FPhaS confidence and high-gradient reliability maps in the same cell field, showing where supported inference succeeds and where high-error pixels remain outside the support claim.

**Supplementary Fig. S5 | Mask-locked FLIM-derived ORR mechanism case.**

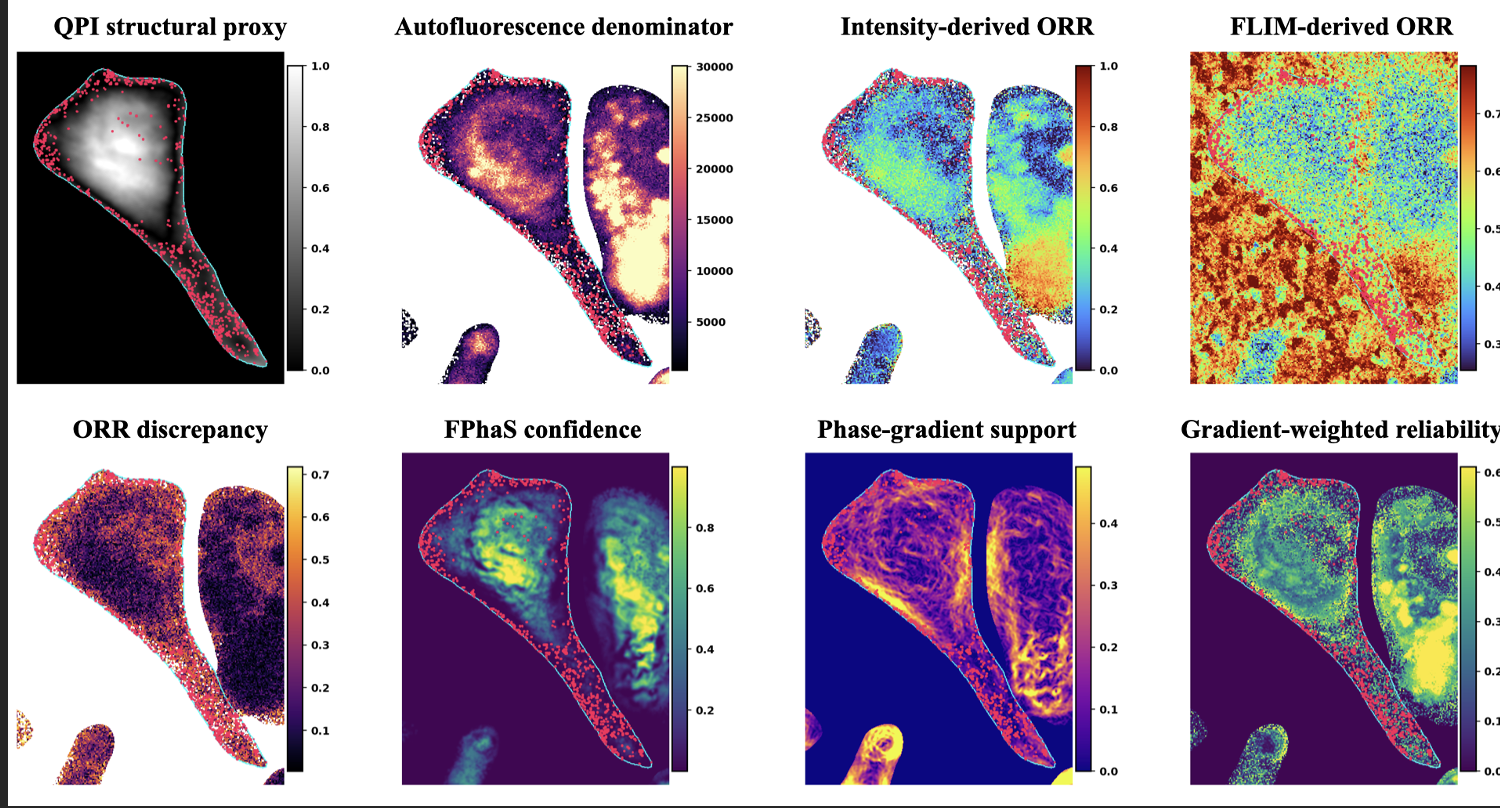

Representative cell-field mechanism case showing QPI dry-mass proxy, SLAM denominator support, intensity ORR, FLIM-derived ORR, absolute intensity-versus-FLIM error, FPhaS QPI-SLAM confidence, high phase-gradient basis and learned/high-gradient reliability. Red markers denote the highest-error pixels and cyan outlines denote cell masks. FLIM-derived ORR is held out as a validation endpoint while FPhaS support maps define where intensity-based inference is reliable.

#### Supplementary Result 6. MCI, MOI and MRI retain support information during aggregation

Supplementary Fig. S6 expands the descriptor analysis behind main Fig. 4. The primary descriptor distributions, descriptor loading structure, cluster-aware landscape, mechanism quadrants and high-end spatial storyboard show how metabolic content, metabolic-structural organization and measurement reliability are retained during pixel-to-cell aggregation. The descriptor cohort is reported separately from the full validation cohort to avoid conflating analysis subsets.

**Supplementary Fig. S6 | MCI, MOI and MRI descriptor robustness.**

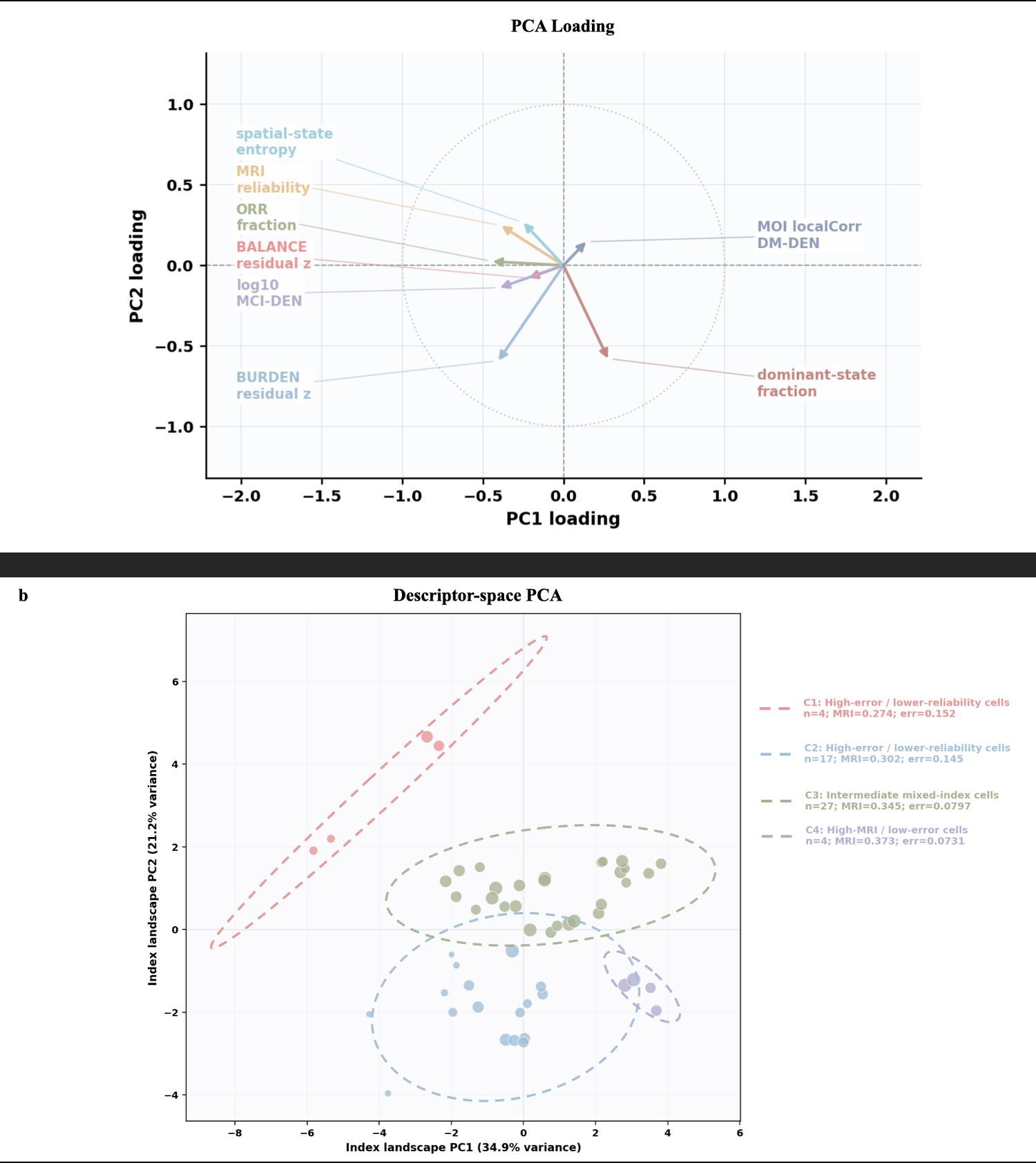

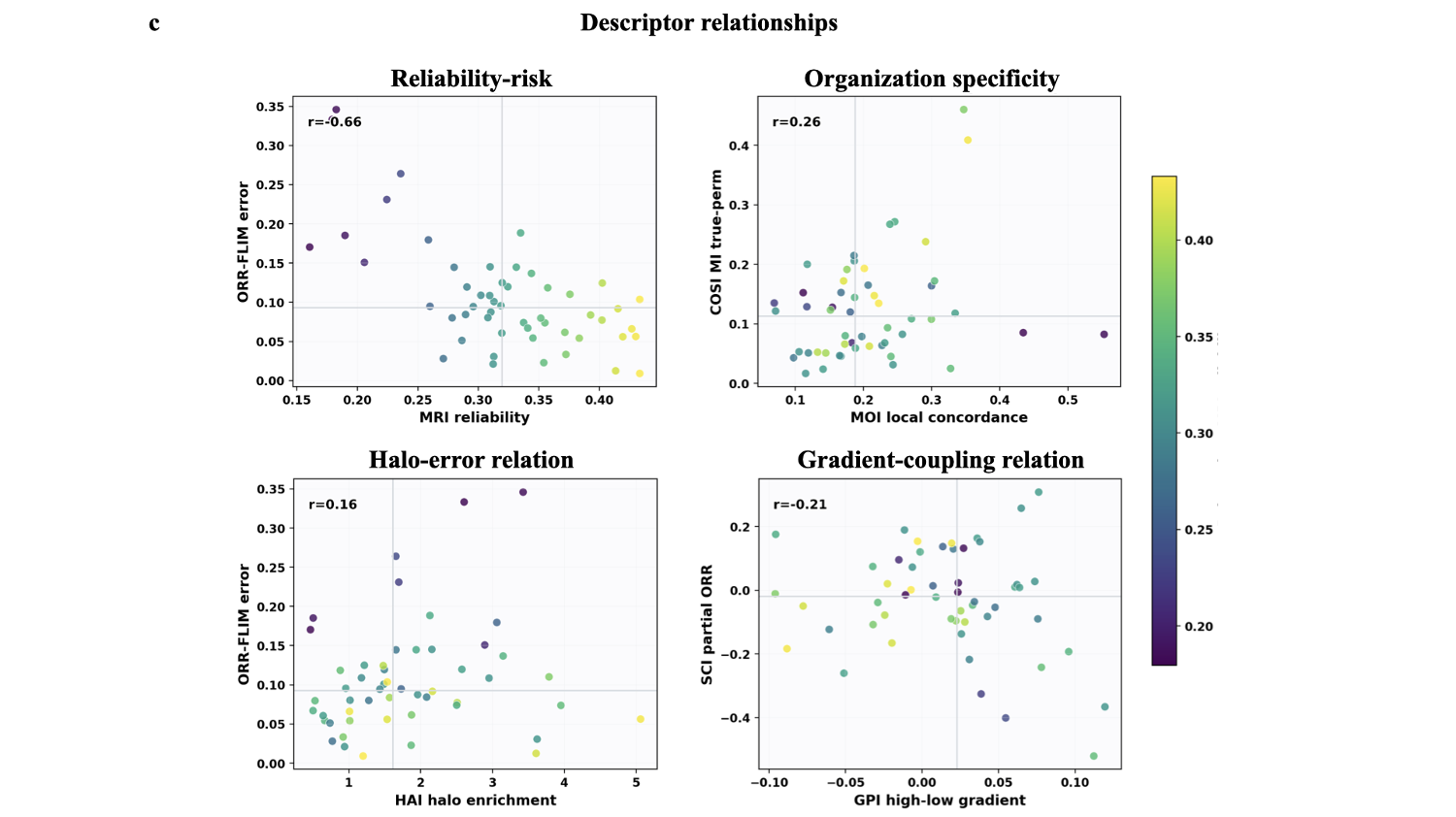

a, show primary descriptor distributions, descriptor PCA loading structure, b, shows cluster-aware primary-cell index landscape, mechanism-quadrant analysis and a high-end spatial storyboard. c, show that MCI, MOI and MRI retain metabolic burden, metabolic-structural organization and measurement-reliability information after cell-level aggregation.

#### Supplementary Result 7. Descriptor-derived states are reproducible image-derived organizations

Supplementary Fig. S7 expands the descriptor-state analysis behind main Fig. 5. PC spatial-score maps, UMAP-to-RGB latent quality-control plots and descriptor-space PCA/UMAP^27^ embeddings support the interpretation that the state map is an image-derived organization of the sampled fields, not a fixed biological taxonomy^28^.

**Supplementary Fig. S7 | Descriptor-state atlas and latent-space quality controls.**

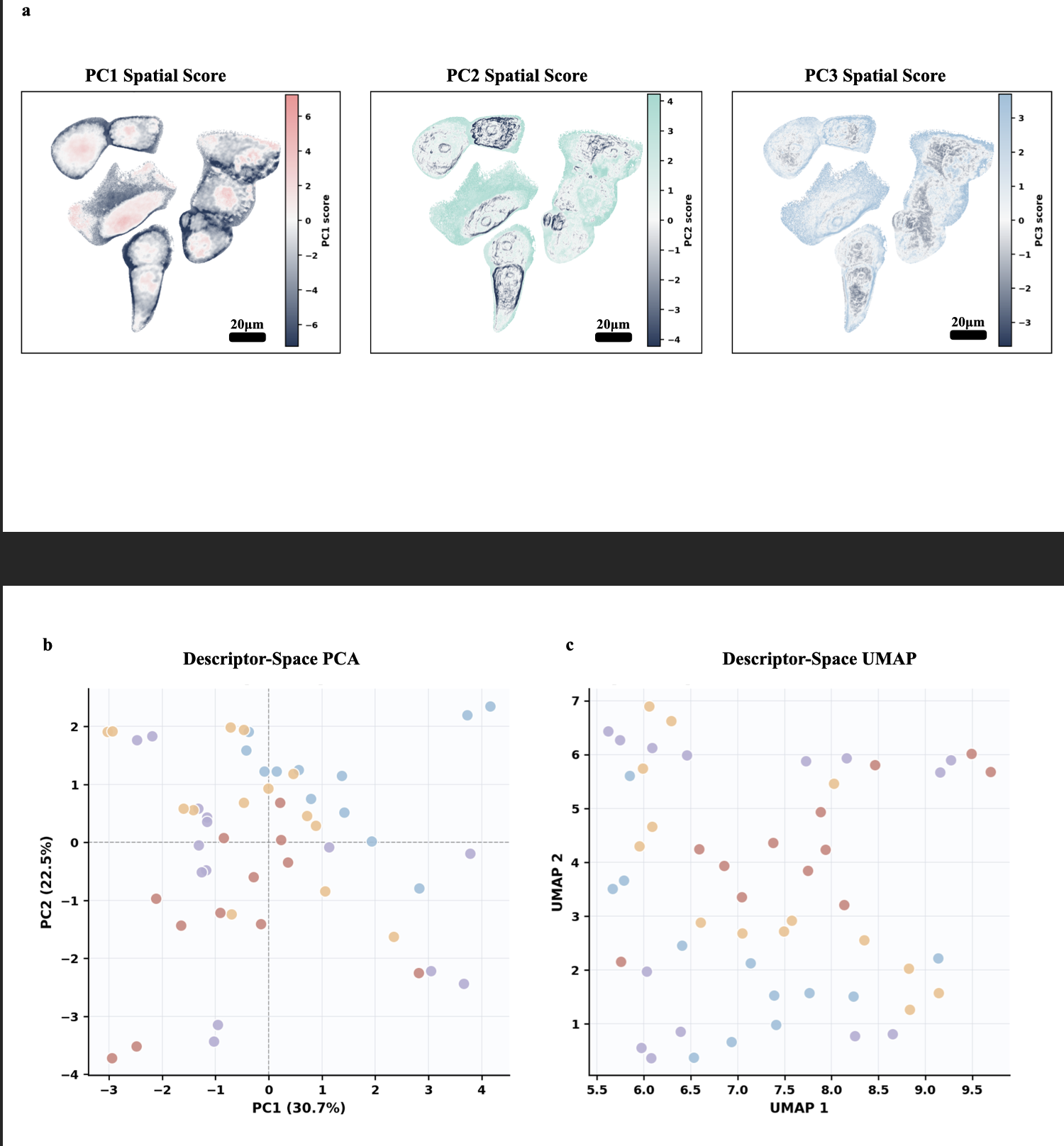

a, show spatial maps of the first three descriptor principal-component scores, b,c, show that descriptor-space PCA/UMAP embeddings of cell-level descriptor measurements. These analyses support interpretation of descriptor-derived states as image-derived organizations of the sampled fields rather than fixed biological classes.

#### Supplementary Result 8. QPI-guided reconstruction and reduced-acquisition benchmarking

Supplementary Figs. S8 and S9 expand the evidence for reconstruction. Supplementary Fig. S8 shows detailed NAD(P)H and FAD reconstruction examples for the reduced input, QPI guide, guided filtering, CombinedWLS and oracle outputs. Supplementary Fig. S9 summarizes file-level paired metric changes from the reduced-acquisition input to CombinedWLS. Together, these controls support a bounded-reconstruction claim: QPI acts as an edge-aware regularizer within the declared support layer, not as an independent source of metabolic contrast.

**Fig. S8 | QPI-guided reconstruction examples and guide-use comparison.**

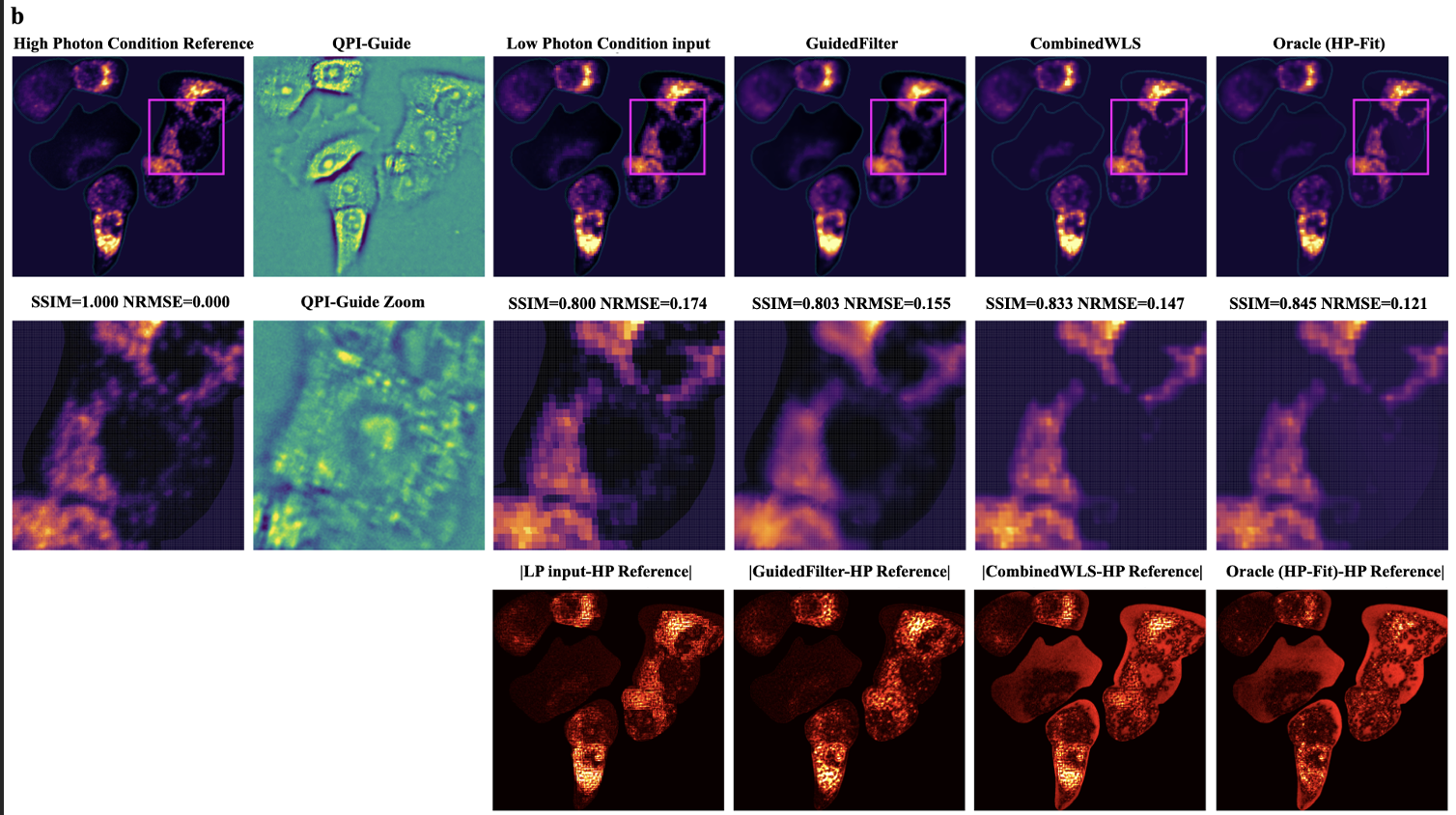

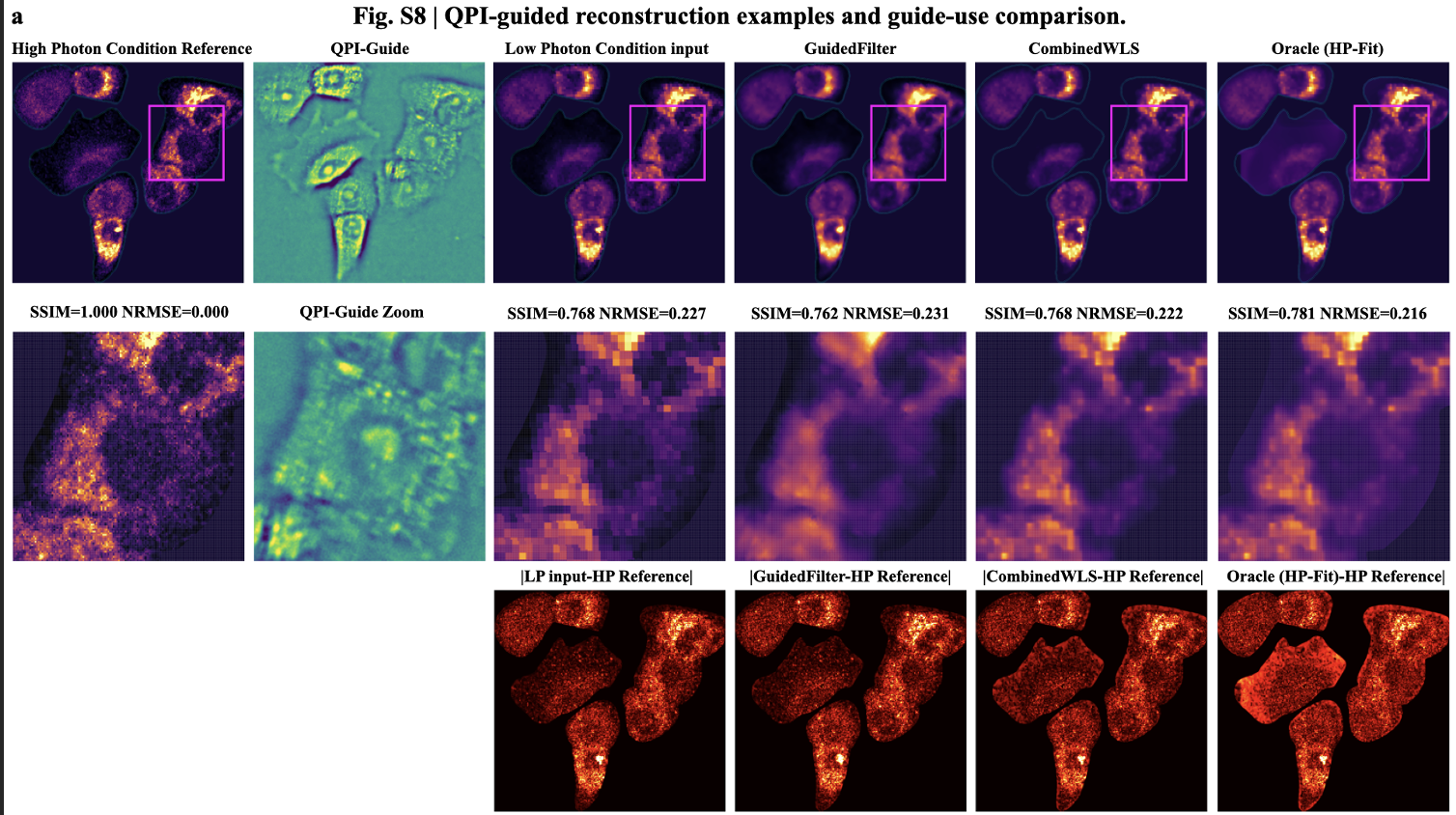

a,b, representative NAD(P)H and FAD reduced-acquisition reconstruction examples show the full reference, QPI phase guide, low-resolution input, guided filtering, CombinedWLS and oracle outputs, with zoomed regions and absolute-difference maps. These panels show that CombinedWLS uses QPI as an edge-aware regularizer while preserving measured fluorescence-derived readouts within the support layer.

**Fig. S9 | Reduced-acquisition metric preservation across imaging units.**

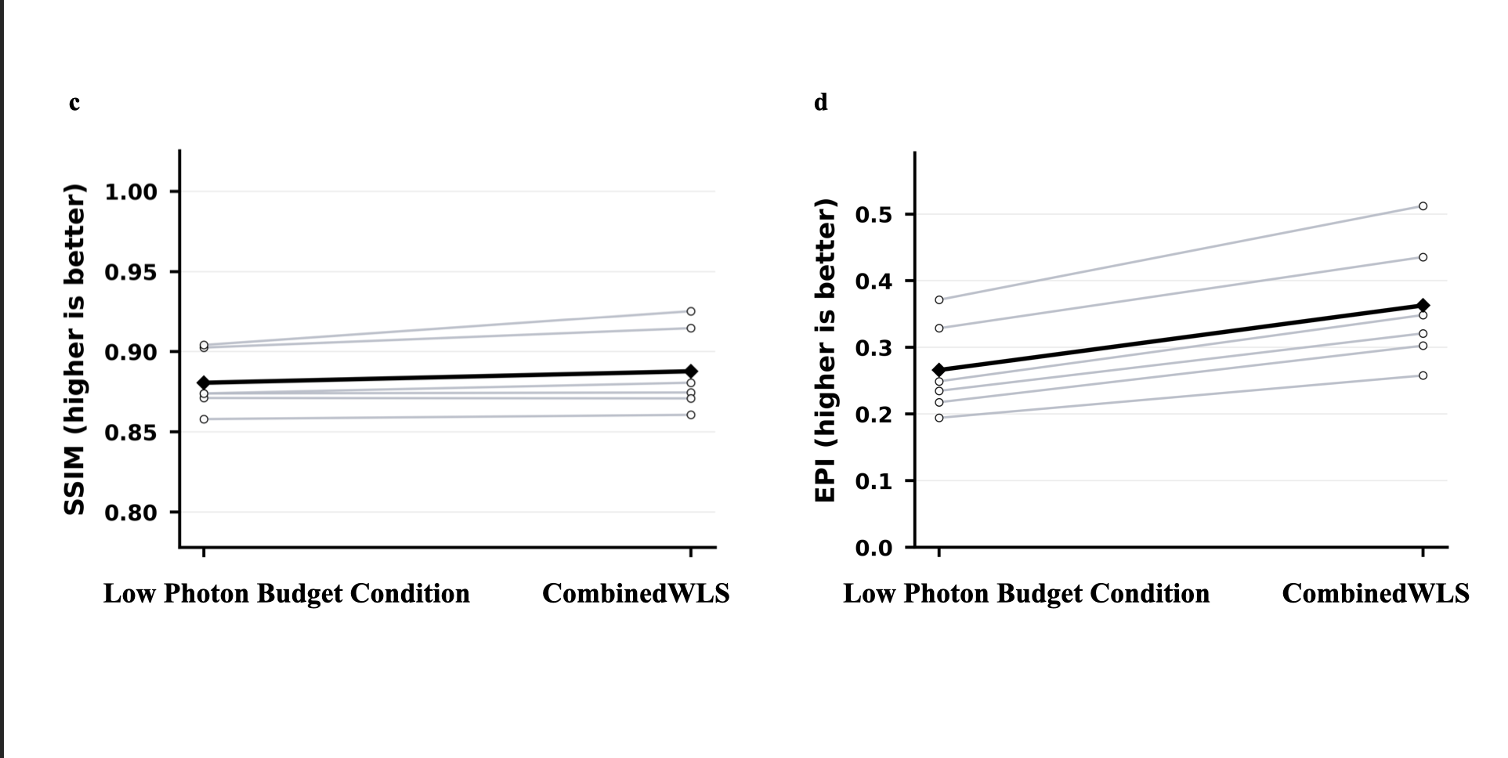

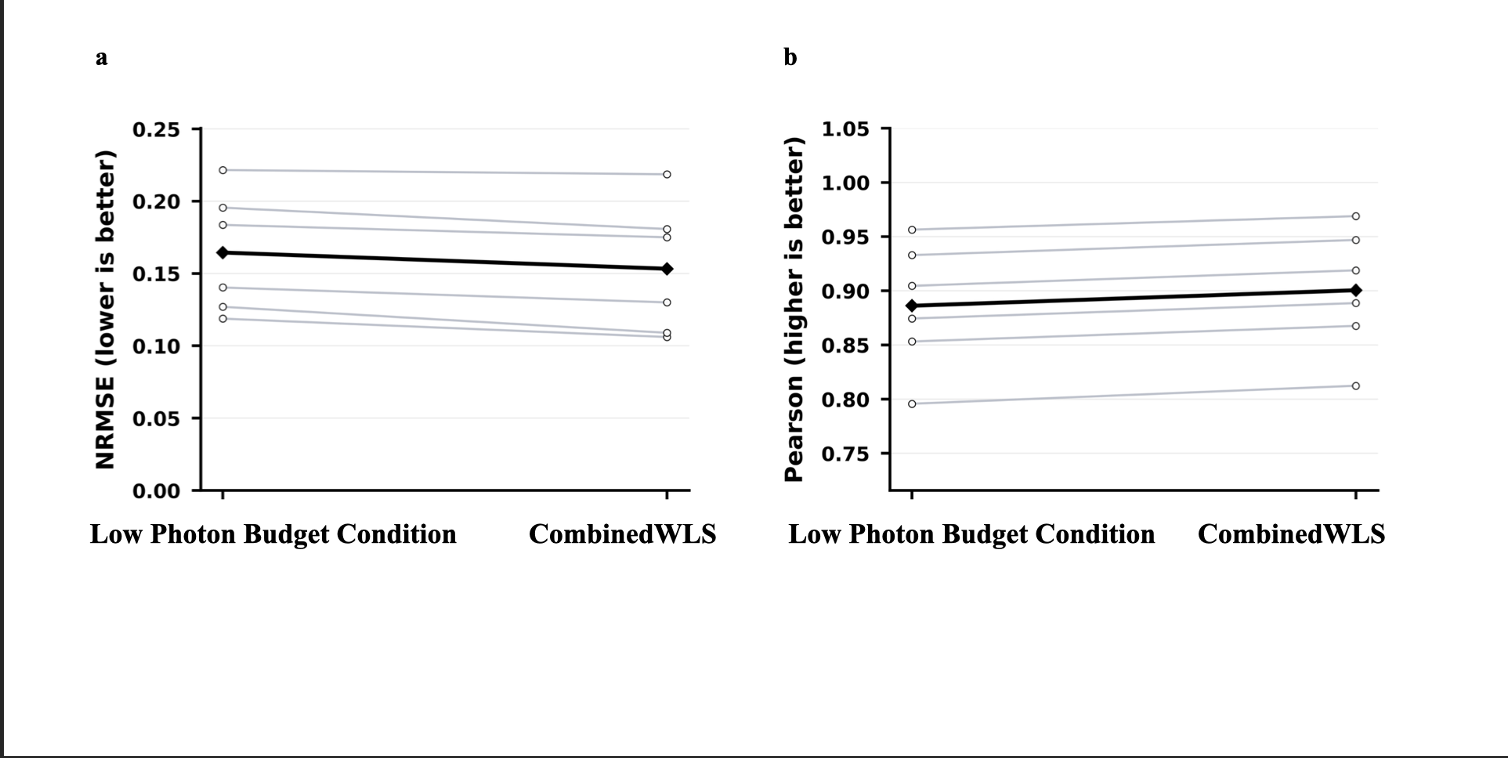

File-level paired metrics compare the reduced-acquisition input with the CombinedWLS reconstruction. a, NRMSE; b, Pearson correlation; c, SSIM; and d, edge-preservation index. Lower is better for NRMSE, and higher is better for Pearson, SSIM, and EPI. Gray lines show paired imaging units, and black summary lines show the aggregate trend.

### Supplementary Tables

Table S1. Multimodal acquisition parameters.

| **Parameter** | **Value** | **Notes** |
| --- | --- | --- |
| Objective | Nikon CFI75 LWD 20XW, 20x, NA 0.8 | Shared objective for QPI, SLAM and FLIM |
| QPI illumination | LED-array DPC, 600 +/- 30 nm | Adafruit 64 x 64 RGB LED matrix |
| SLAM excitation | 1000-1150 nm femtosecond excitation | Average sample-plane power 30 mW |
| NAD(P)H channel | 450 +/- 30 nm | Autofluorescence channel |
| FAD channel | 580 +/- 30 nm | Autofluorescence channel |
| SHG / THG | 515 +/- 20 nm / 385 +/- 20 nm | Harmonic-generation channels |
| SLAM sampling | 512 x 512 over 100 micrometre | 0.195 micrometre per pixel |
| QPI sampling | 490 micrometre field sampled at 0.109 micrometre per pixel | Registered to SLAM field |
| FLIM | PicoHarp 330, 8 ps TCSPC bin | Lifetime-resolved biochemical contrast; held out from confidence-map construction |

| **Component** | **Definition** | **Purpose** |
| --- | --- | --- |
| Mass support | Robustly scaled smoothed QPI mass-support image | Identifies phase-supported cellular material |
| Interior support | Distance-to-boundary term with d0 = 6 pixels | Reduces boundary mixing |
| Low-gradient support | One minus robustly scaled QPI gradient magnitude | Downweights sharp boundary/edge ambiguity |
| Low-texture support | One minus robustly scaled local QPI texture | Downweights local structural heterogeneity that may reduce ratio stability |

Table S2. QPI structural reliability components.

Table S3. Descriptor outputs to report from source data.

| **Output** | **Value** | **Interpretation** |
| --- | --- | --- |
| Analyzed cells | 72 cells from the descriptor subset; validation cohort reported separately as 10 fields of view across 5 samples | Descriptor cohort, not the full validation cohort |
| MCI_DEN median | 22,573.5 | Support-normalized burden descriptor |
| MOI local correlation median | 0.188 | Local metabolic-structural organization |
| MRI median | 0.319 | Reliability descriptor |
| Between-condition CV | 0.0537 | Descriptor stability under reduced photon support |
| Bootstrap ARI | 0.991 | State stability |
| Cross-FOV ARI | 0.339 | Transfer across fields |
| Cramer V | 0.147 | Modest redox association |

| **Output** | **Value or required entry** | **Main use** |
| --- | --- | --- |
| CombinedWLS support-layer NRMSE | 0.1532 | Primary reconstruction accuracy |
| CombinedWLS support-layer Pearson | 0.901 | Intensity/structure agreement |
| CombinedWLS support-layer SSIM | 0.888 | Structural similarity |
| CombinedWLS support-layer EPI | 0.363 | Edge preservation |
| Oracle recovery | 45.3% NRMSE; 91.4% Pearson; 51.0% SSIM; 99.9% EPI | Recovery relative to oracle bound |
| Reduced-acquisition factor | 8x | Acquisition-burden claim |
| Reduced-acquisition source mapping | Full-acquisition reference and eightfold-reduced fluorescence-acquisition condition | Maps reduced-acquisition analysis to final manuscript labels |

Table S4. Reconstruction and reduced-acquisition outputs to report from source data.

### Supplementary References

1.	Tian, L. & Waller, L. Quantitative differential phase contrast imaging in an LED array microscope. *Opt. Express* **23**, 11394–11403 (2015).

2.	Ku, H. H. Notes on the use of propagation of error formulas.

3.	Oehlert, G. W. A Note on the Delta Method. *Am. Stat.* **46**, 27–29 (1992).

4.	Muir, R. D., Kissick, D. J. & Simpson, G. J. Statistical connection of binomial photon counting and photon averaging in high dynamic range beam-scanning microscopy. *Opt. Express* **20**, 10406–10415 (2012).

5.	Georgakoudi, I. *et al.* Consensus guidelines for cellular label-free optical metabolic imaging: ensuring accuracy and reproducibility in metabolic profiling. *J. Biomed. Opt.* **30**, S23901 (2025).

6.	Park, Y., Depeursinge, C. & Popescu, G. Quantitative phase imaging in biomedicine. *Nat. Photonics* **12**, 578–589 (2018).

7.	Popescu, G. *et al.* Optical imaging of cell mass and growth dynamics. *Am. J. Physiol.-Cell Physiol.* **295**, C538–C544 (2008).

8.	Phipson, B. & Smyth, G. K. Permutation P-values Should Never Be Zero: Calculating Exact P-values When Permutations Are Randomly Drawn. *Stat. Appl. Genet. Mol. Biol.* **9**,.

9.	He, K., Sun, J. & Tang, X. Guided Image Filtering. *IEEE Trans. Pattern Anal. Mach. Intell.* **35**, 1397–1409 (2013).

10.	Huber, P. J. Robust Estimation of a Location Parameter. *Ann Math Stat.* **35**, 73–101 (1964).

11.	Hestenes, M. R. & Stiefel, E. Methods of conjugate gradients for solving linear systems.

12.	Wang, Z., Bovik, A. C., Sheikh, H. R. & Simoncelli, E. P. Image quality assessment: from error visibility to structural similarity. *IEEE Trans. Image Process.* **13**, 600–612 (2004).

13.	Pedregosa, F. *et al.* Scikit-learn: Machine Learning in Python. *Mach. Learn. PYTHON*.

14.	Efron, B. Bootstrap Methods: Another Look at the Jackknife. *Ann. Stat.* **7**, 1–26 (1979).

15.	Datta, R., Heaster, T. M., Sharick, J. T., Gillette, A. A. & Skala, M. C. Fluorescence lifetime imaging microscopy: fundamentals and advances in instrumentation, analysis, and applications. *J. Biomed. Opt.* **25**, 071203 (2020).

16.	Ranjit, S., Malacrida, L., Jameson, D. M. & Gratton, E. Fit-free analysis of fluorescence lifetime imaging data using the phasor approach. *Nat. Protoc.* **13**, 1979–2004 (2018).

17.	Lakowicz, J. R., Szmacinski, H., Nowaczyk, K. & Johnson, M. L. Fluorescence lifetime imaging of free and protein-bound NADH. *Proc. Natl. Acad. Sci.* **89**, 1271–1275 (1992).

18.	Chance, B. Optical method. *Annu. Rev. Biophys. Biophys. Chem.* **20**, 1–30 (1991).

19.	Skala, M. C. *et al.* In vivo multiphoton microscopy of NADH and FAD redox states, fluorescence lifetimes, and cellular morphology in precancerous epithelia. *Proc. Natl. Acad. Sci.* **104**, 19494–19499 (2007).

20.	Fitzpatrick, J. M., West, J. B. & Maurer, C. R. Predicting error in rigid-body point-based registration. *IEEE Trans. Med. Imaging* **17**, 694–702 (1998).

21.	Modersitzki, J. *Numerical Methods for Image Registration*. (OUP Oxford, 2003).

22.	Baba, K., Shibata, R. & Sibuya, M. Partial Correlation and Conditional Correlation as Measures of Conditional Independence. *Aust. N. Z. J. Stat.* **46**, 657–664 (2004).

23.	Pearson, K. LIII. On lines and planes of closest fit to systems of points in space. *Lond. Edinb. Dublin Philos. Mag. J. Sci.* **2**, 559–572 (1901).

24.	You, S. *et al.* Label-free deep profiling of the tumor microenvironment. *Cancer Res.* **81**, 2534–2544 (2021).

25.	Boppart, S. A., You, S., Li, L., Chen, J. & Tu, H. Simultaneous label-free autofluorescence-multiharmonic microscopy and beyond. *APL Photonics* **4**, 100901 (2019).

26.	Boppart, S. A. *et al.* Label-free optical imaging technologies for rapid translation and use during intraoperative surgical and tumor margin assessment. *J. Biomed. Opt.* **23**, 021104 (2017).

27.	McInnes, L., Healy, J., Saul, N. & Großberger, L. UMAP: Uniform Manifold Approximation and Projection. *J. Open Source Softw.* **3**, 861 (2018).

28.	Hubert, L. & Arabie, P. Comparing partitions. *J. Classif.* **2**, 193–218 (1985).
